# One-Pot Enzymatic ADDing of Click Chemistry Handles for Protein Immobilization and Bioconjugation of Small and Biomolecules

**DOI:** 10.1101/2025.05.17.650163

**Authors:** Wahyu S. Widodo, Xiaoman Fan, Maximilian J.L.J Fürst

**Affiliations:** Molecular Enzymology Group, University of Groningen, Nijenborgh 3, Groningen, 9747AG, The Netherlands

## Abstract

Site-specific attachment of biorthogonal handles to proteins is an essential tool in chemical biology research and diverse applications including imaging and protein immobilization, as well as for the development of next-generation therapeutics such as antibody drug-conjugates. Among the available methods, enzymatic post-translational modification of short protein tags offers precision, stability, and modularity. However, broader application is often limited by complex substrate syntheses, the requirement of long or rigid recognition tags, and limited reaction efficiencies. Here, we present ADDing, a straightforward enzymatic method for functionalizing proteins with click chemistry handles using the flavin transferase ApbE. We discovered that, given a dedicated adenine diphosphate derivative (ADD) substrate, the enzyme attaches a phosphoribosyl moiety bearing bioorthogonal handles to proteins featuring a DxxxGAT amino acid motif. As the substrates can easily be enzymatically synthesized from NAD and inexpensive precursors, ADDing click handles can be performed in a streamlined, one-pot workflow combining substrate synthesis and protein conjugation. ADDing allows rapid, high-yield functionalization of proteins featuring the recognition tag at either terminus or internal loops and is compatible with copper and copper free azide-alkyne cycloaddition reactions. To demonstrate its broad applicability, we performed a wide variety of protein functionalizations, including fluorescent labeling, protein-protein, protein-DNA conjugation, and protein immobilization. This versatile technology thus holds great potential for chemical biology and the production of biological therapeutics.

## Introduction

Molecular attachment to biomolecules is a fundamental technique in chemical biology that enables functionalization, tracking, and manipulation of proteins and nucleic acids for applications in imaging, immobilization, and mechanistic studies.^[1–3]^ Precise conjugation strategies are furthermore crucial for next-generation therapeutics such as antibody-drug conjugates and modified peptides, which enable controlled bioactive compound delivery.^[4–7]^ Moreover, side-chain decoration, branched polypeptides, and protein-nucleic acid conjugates are not only of great interest for biomedical and biotechnological applications, but provide essential tools for the study of post-translational modifications (PTMs) and cellular processes such as protein degradation.^[8–10]^

Covalent protein modifications can be achieved through stochastic conjugation or site-specifically. Stochastic methods exploit the inherent reactivity of some amino acids, typically targeting cysteine or lysine residues.^[11]^ While applicable to native proteins, this approach often leads to uncontrolled multivalency or requires mutagenesis for monovalent attachment. Additionally, random modifications can disrupt protein function, and the chemical reaction may be detrimental to protein stability or incompatible with the desired application.^[12,13]^ To overcome these limitations, biochemical strategies for site-specific modification have been developed, including non-canonical amino acid (ncAA) incorporation and enzymatic post-translational modifications (PTMs).^[11,14,15]^ ncAAs are introduced by hijacking the translation process via engineered tRNA and aminoacyl-tRNA synthetases, and provide bioorthogonal handles as amino acids with unique reactivity. Although well-established, this approach requires complex expression systems and can result in low incorporation efficiency and heterogeneous products.^[16]^ In contrast, enzymatic PTMs offer an efficient and scalable alternative, without interfering in translation.^[11]^ These methods employ promiscuous enzymes that target specific peptide tags, enabling precise, stable, and modular protein modifications.^[15,17]^ When genetic manipulation of the target is feasible, PTM strategies are thus generally the preferred method for protein labelling.^[15]^ Regardless of the enzyme used, protein labelling typically follows two steps: 1) enzymatic conjugation of a reactive handle and 2) chemical attachment of the desired functional molecule. This approach reduces synthetic complexity and accounts for the limits in promiscuity of the modifying enzymes. To maximise yields, minimize system-specific adjustments, and prevent side products, the chemical step must be biocompatible and orthogonal. A breakthrough in this regard was the advent of click chemistry, which enabled highly selective biocompatible reactions, significantly expanding the toolkit for biomolecular modifications and gaining widespread adoption.^[18,19]^

The various enzymatic PTM systems that were developed differ in their acceptor tag requirements and the necessary chemistry on the donor molecule. While the optimal choice is application-dependent, short peptide tags and synthetically accessible substrates are highly preferred. A recent review identified eight enzyme systems capable of attaching orthogonal handles such as azides, alkynes, and tetrazines to peptide tags.^[11,15]^ Examples include transferases such as farnesyltransferase, which recognizes a C-terminal CaaX motif^[20]^, and ligases,^[21]^ such as tyrosine (TTL)^[22–25]^ or biotin ligase (BirA)^[26,27]^ and sortase.^[28]^ Despite significant progress, the search continues for an ideal enzyme that combines easily synthesizable substrates, a short and flexible recognition peptide, and high activity.^[29–34]^

Inspired by studies demonstrating the use of the flavin transferase ApbE for site-specific attachment of flavin derivates to tag proteins,^[35,36]^ we here describe ADD-tagging or ADDing—a system for incorporating click chemistry handles using enzymatically synthesized substrates transferred via ApbE. In nature, ApbE attaches FMN to a threonine or serine in the B and C subunits of the bacterial respiratory sodium pump Nqr via a conserved DxxxGA[T/S] motif.^[37–39]^ Based on the reported substrate preference in the so-called flavin tag,^[35,36]^ we hypothesized that ApbE is active on other dinucleotides, recognizing an AMP moiety and transferring the remaining portion to target proteins. By leveraging an enzymatic strategy via ADP-ribosyl cyclase (ADPRC)^[40]^ for the synthesis of dinucleotides featuring a click chemistry handle and subsequent ApbE-enabled conjugation, we demonstrate the successful decoration of proteins with small molecules such as biotin and fluorophores, as well as supramolecular structures like DNA, proteins, and beads (**Scheme 1**). ADDing enables substrate synthesis and ApbE-mediated conjugation in a streamlined one-pot process, simplifying the workflow and eliminating the need for intermediate purification. Our system thus establishes a valuable alternative for diverse chemical biology applications including decoration with small and biomolecules, as well as protein immobilization.

**Scheme 1.**
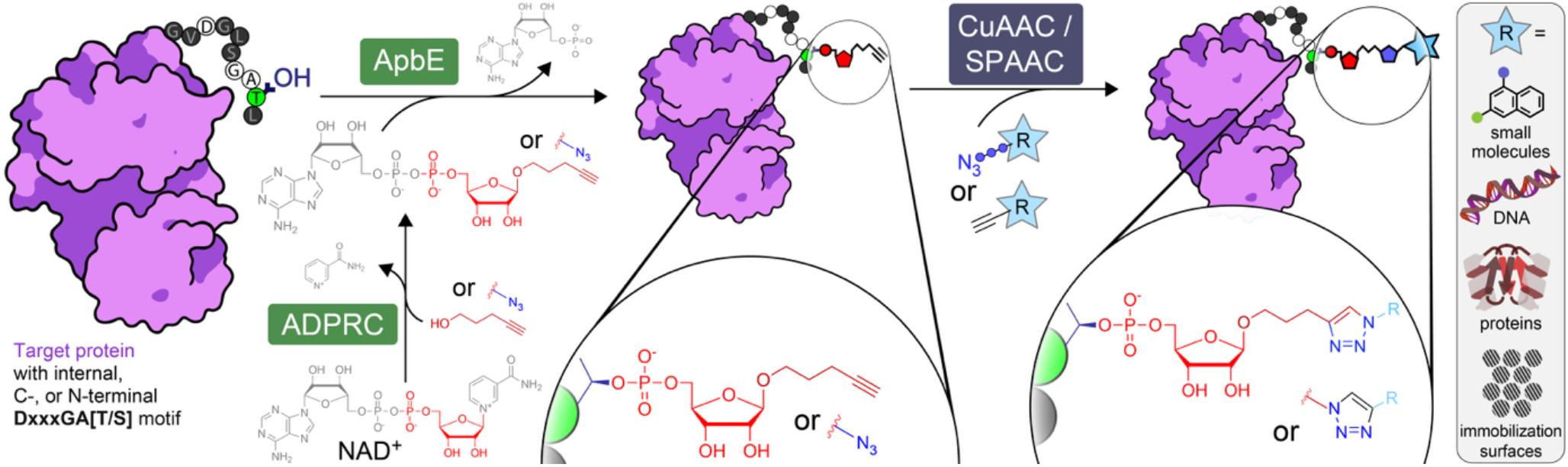
Overview of ADD-tagging including ADP-ribosyl cyclase (ADPRC)-catalyzed dinucleotide substrate generation and two step chemoenzymatic labelling of target proteins with the flavin transferase ApbE, followed by click chemistry-based functional group attachment.

## Results and Discussion

We began our study with the premise that ApbE, previously reported to accept not only its natural substrate, flavin adenine dinucleotide (FAD),^[35]^ but also the structurally distinct NADH,^[36]^ exhibits broad substrate specificity. This flexibility suggested the potential to exploit ApbE for attaching other functional groups. Given that both FAD and NADH share a dinucleotide structure, and that AMP is consistently released as a byproduct during catalysis, we hypothesized that ApbE could also accommodate other AMP-based dinucleotides, including synthetic variants. While chemical synthesis of dinucleotides is well established,^[41]^ we were intrigued by the possibility of achieving protein modification via a synthesis-free, enzyme-only route. Specifically, we considered using adenine diphosphate ribose cyclase (ADPRC) to generate the required precursor.^[40]^ In nature, certain eukaryotic phyla produce ADPRC to catalyze the formation of the signaling molecule cyclic ADP-ribose from NAD^+^. However, it was recently discovered that its enzymatic mechanism can be redirected to an alternative trans glycosylation reaction, where the nicotinamide moiety is replaced by a nucleophilic donor molecule,^[42]^ including small alkynyl and azido alcohols.^[43,44]^ Recognizing the outstanding utility of click-chemistry handles for bioconjugation, we devised the following strategy: (i) enzymatic synthesis of click-chemistry-compatible dinucleotides from NAD using ADPRC, (ii) ApbE-mediated coupling to a target protein bearing the ApbE recognition tag, and (iii) subsequent attachment of a click-chemistry-compatible molecule.

### Producing apo ApbE

While ADPRC can readily be obtained commercially, recombinant ApbE production in *E. coli* required some consideration. Overexpression in standard *E. coli* strains such as BL21 yielded a holoenzyme in complex with FAD, evident from its yellow color **(Figure S1)**. As the residual FAD led to undesired flavinylated target protein in reactions without additional flavin (**Figure S2**), we produced apo-ApbE in a dedicated riboflavin-deficient *E. coli* BL21(AI) strain with a deletion in the riboflavin biosynthesis *ribB* (manuscript in preparation). Spectrophotometry (**Figure S1**) and control reactions analyzed via MS confirmed the so-obtained ApbE to be in the apo form **(Figure S2a and S2c)**.

### Enzymatic synthesis of pent-4-yn-1-yl ADP-ribose and 3-azido-propyl ADP-ribose

Next, we performed reactions with commercial ADPRC from *Aplysia californica* using NAD^+^ and either 4-pentyn-1-ol or 3-azido propanol, following an established procedure^[43,44]^ (**Figure 1a**). Ultra-high-performance liquid chromatography (UHPLC) confirmed the conversion of NAD^+^ into alkyne- and azide-functionalized adenosine diphosphate ribose (ADPR). We refer to the resulting products, pent-4-yn-1-yl ADPR and 3-azido-propyl ADPR, hereafter as alkynyl-ADPR and azido-ADPR. Our attempts to reduce the relatively high alcohol substrate concentrations were unsuccessful, confirming the need for a high (> 7-fold) excess to achieve trans glycosylation in good yields (**Figure S3**). Using 770 mM alcohol, the main peaks observed in HPLC analyses corresponded to the azido and alkynyl ADPRs (**Figure 1b**). Noticing minor amounts of cyclic cADPR product, we tested a potential inhibitory effect of this molecule on ApbE, but the enzyme showed neither activity nor inhibition with either cADPR or NAD^+^ **(Figure S4)**. The inactivity with the oxidized nicotinamide cofactor was unexpected, suggesting that ApbE promiscuity is constrained by certain molecular features, possibly the positively charged group. Interestingly, this behavior prevents a potential interference of unreacted starting material and enables a seamless tandem use of the two enzymatic steps. Thus, we deemed the ADPRC bioconversions suitable for the subsequent ApbE reaction without further purification.

**Figure 1.**
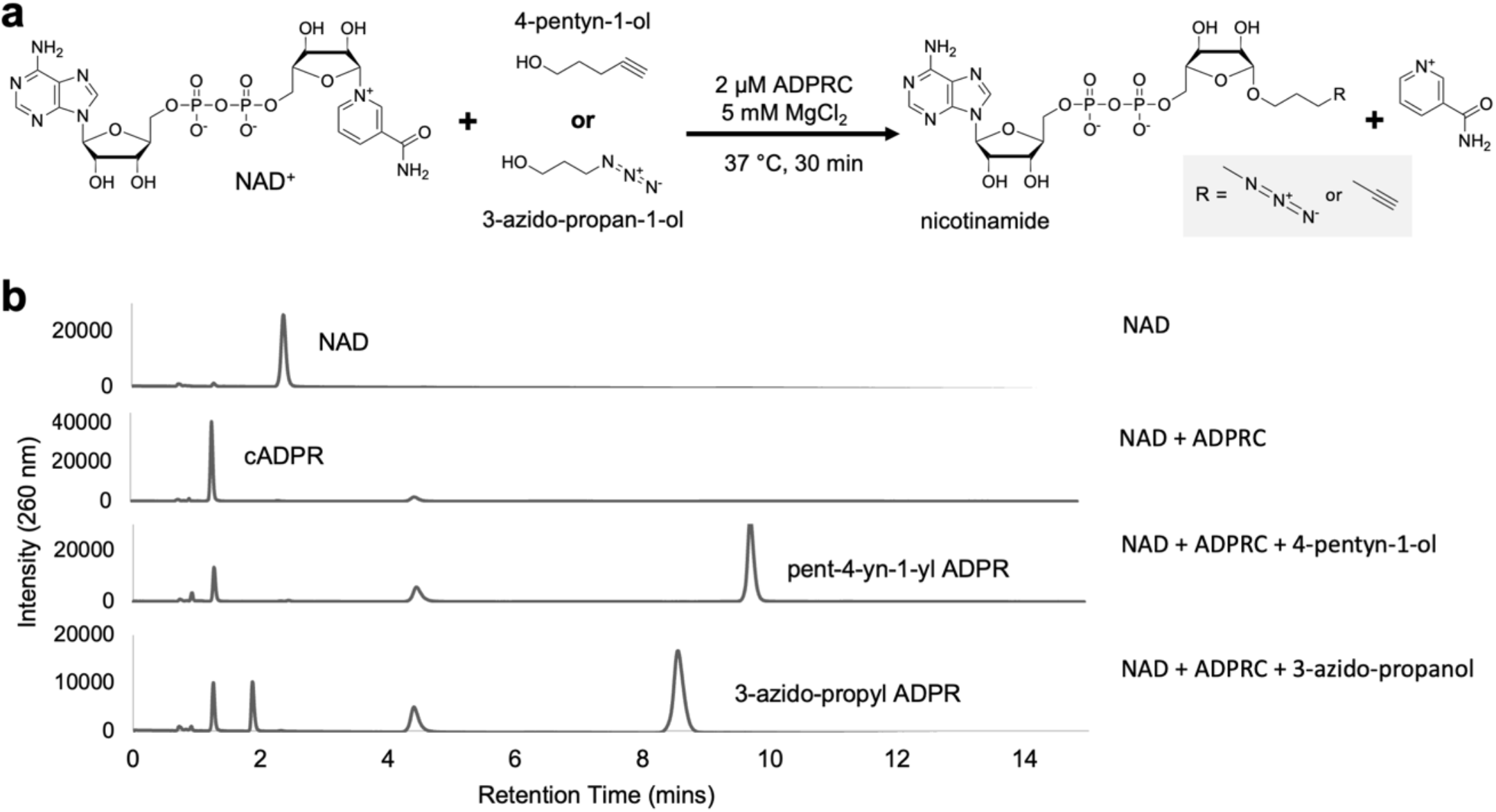
ADPRC-catalyzed trans-glycosylation reactions. a) Schematic representation of the ADPRC reaction illustrating the generation of alkynyl and azido ADPR. b) UHPLC chromatograms of ADPR reactions with 4-penyn-1-ol and 3-azido-propanol and controls.

**Figure 2.**
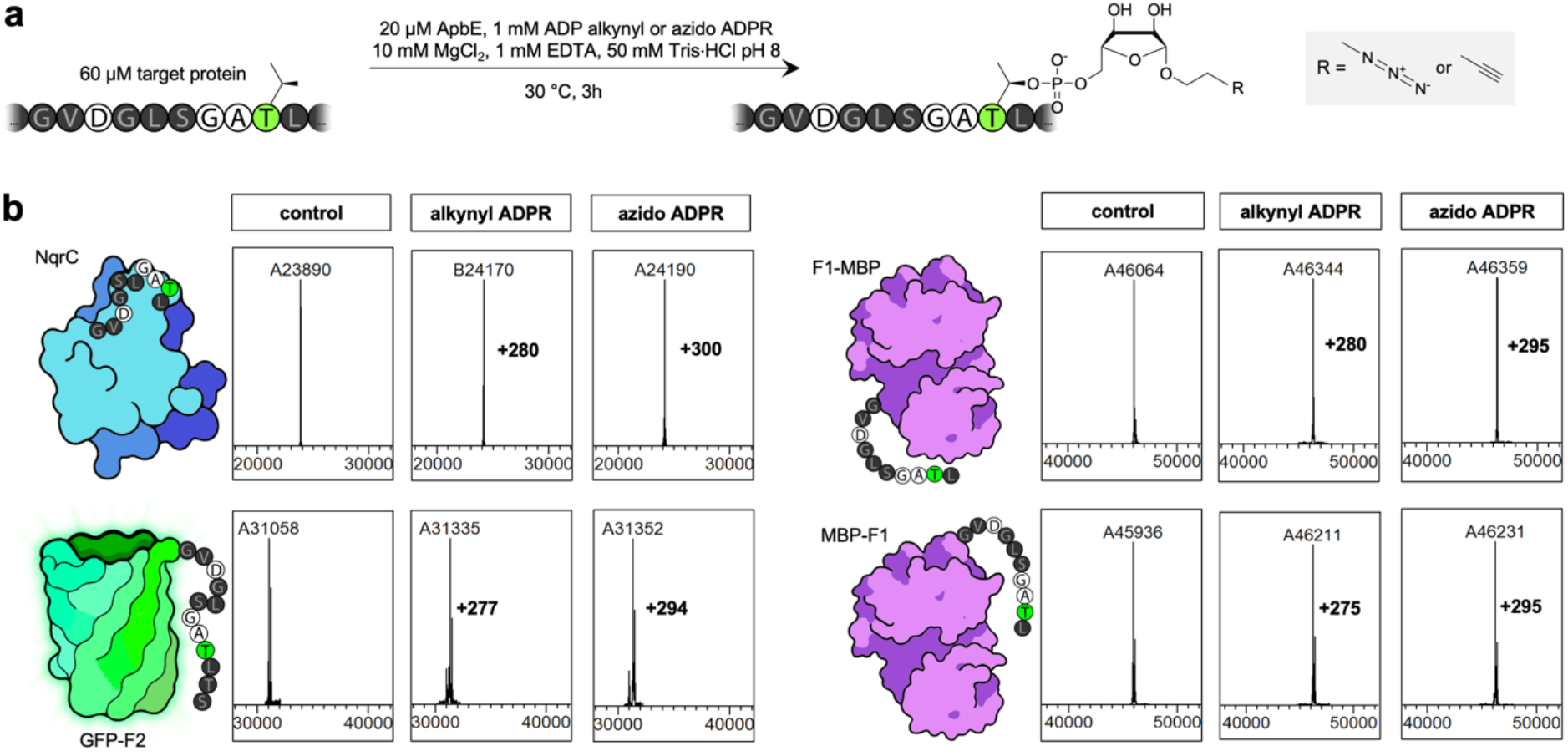
Bioorthogonal handles (alkynyl and azido ADPR) conjugation to various proteins with ADD-tag located C-terminally, N-terminally and at an internal loop. (a) Reaction conditions for the attachment of bioorthogonal handles to target proteins using ApbE. (b) MS analysis showing the successful incorporation of alkynyl and azido ribose phosphate, indicated by mass additions of 281 Da and 296 Da, respectively. “F1” and “F2” denote the tag sequences GVDGLSGA**T**LTS and GVDGLSGA**T**L, respectively.

### Attachment of alkynyl and azido ribose phosphate on proteins

We then moved to investigate whether ApbE accepts alkynyl and azido ADPR as donor substrates. Analogous to FMN transfer from FAD, we expected ApbE to attach the alkynyl ribose phosphate and azido ribose phosphate moieties from the modified ADPRs. As the first acceptor protein, we tested NqrC, the enzyme’s natural target. This protein is a subunit of the Na^+^-translocating NADH:quinone oxidoreductase enzyme complex and features the recognition motif at an internal loop-helix junction. Additionally, we produced various recombinant proteins featuring the motif as peptide tags: maltose binding protein (MBP), small ubiquitin-like modified (SUMO), and enhanced green fluorescent protein (eGFP). For flavinylation, ApbE was found to be highly permissive toward the acceptor protein, allowing targets to bear either an optimized (“F1 tag”) or shortened peptide tag (“F2”) at termini or internal loops.^[35]^ For detection of the protein modifications, we employed electrospray ionization mass spectrometry analysis (ESI-MS).

Gratifyingly, we found that reactions under the previously described standard conditions (1 mg/mL ApbE, 3 mg/mL target proteins, 3 h, 30 °C) resulted in the complete modification of NqrC, MBP-F1 (C-terminal F1), and F1-MBP (N-terminal F1) (**Figure 1b**), as evidenced by a complete mass shift in deconvoluted mass spectrometry (MS) profiles. GFP-F2 displayed slightly reduced labelling kinetics, requiring an extended incubation of 12 hours to achieve complete conversion. Only one of our test constructs, SUMO-iF1 (internal F1 tag), led to incomplete conversion after 3 hours, and even extended reactions of 12 hours led to only minimal additional modification (**Figure S5**). Having thus established that ApbE is a highly efficient and versatile enzymatic tool to add adenine dinucleotide derived (ADD) chemical moieties to proteins, we named the here established principle ADD-taging or ADDing. Although we employed two previously optimized peptide tags of 12 (F1) and 10 (F2) amino acids,^[35]^ the minimal ApbE recognition motif comprises only seven semi-flexible amino acid (D-[GAIQ]-[IALVF]-[ST]-G-A-[ST]).^[37]^ This places the ADD tag among the shortest peptide tags for labelling,^[15,17]^ especially considering that the shortest recognition tags, such as the tag used for the SUMO-conjugating enzyme Ubc9, in practice requires the addition of amino acid linkers.^[45]^ In comparison, other commonly used tags, such as tubulin tyrosine ligase,^[29,46]^ and lipoic acid ligase,^[15,17]^ are significantly longer. Moreover, ADDtag’s sequence flexibility can facilitate internal insertions of the motif with minimal alterations, as previously demonstrated using computationally-aided design.^[47]^

Given our initial concerns about ApbE copurifying in complex with FAD in standard strains, we also assessed the practical impact of such a scenario. Although in line with our expectations, we indeed found FMNylated protein, the majority of the target proteins incorporated the click chemistry handles when using the same reaction conditions as before (**Figure S2b**), indicating that a dedicated flavin-free strain is not a strict requirement for the production of ApbE for ADDing.

We also evaluated the stability of the phosphodiester linkage between threonine and the attached molecules against enzymatic degradation by various phosphatases and nucleases (**Figure S6**). To facilitate analysis, flavin mononucleotide (FMN) was used as a model molecule attached to NqrC, and the resulting FMNylated protein was visualized by SDS-PAGE, taking advantage of FMN’s intrinsic fluorescence. The attached FMN remained intact after enzymatic treatment, indicating that the ADDing modification provides a stable protein functionalization (**Figure S6**). In addition, this linkage has previously been shown to tolerate a broad range of pH, temperatures, and buffer additives.^[35]^

### One-pot ADPRC and ApbE reactions

Given ApbE’s inactivity towards NAD^+^ and cADPR, we assessed the possibility of substrate generation and conjugation in a single pot. Aiming to establish an ideal compromise in the conditions for both ApbE and ADPRC, we assessed various concentrations of substrates while maintaining a ∼7x excess of alcohol precursor. Since we observed protein precipitations at concentrations exceeding 100 mM of 4-pentyn-1-ol—indicative of a detrimental effect of the alcohol on ApbE and/or target protein—we settled on utilizing 1.5 mM NAD^+^ and 100 mM alcohol. Under these conditions, incorporation of alkynyl phospho-ribose into F1-MBP was successfully completed within 12 hours, with significant protein modification occurring already after 3 hours. Conducting the reaction in Tris buffer pH 8, the standard buffer for ApbE, led to a slightly faster conversion than using the ADPRC-preferred buffer HEPES pH 7 (**Figure 3**).

**Figure 3.**
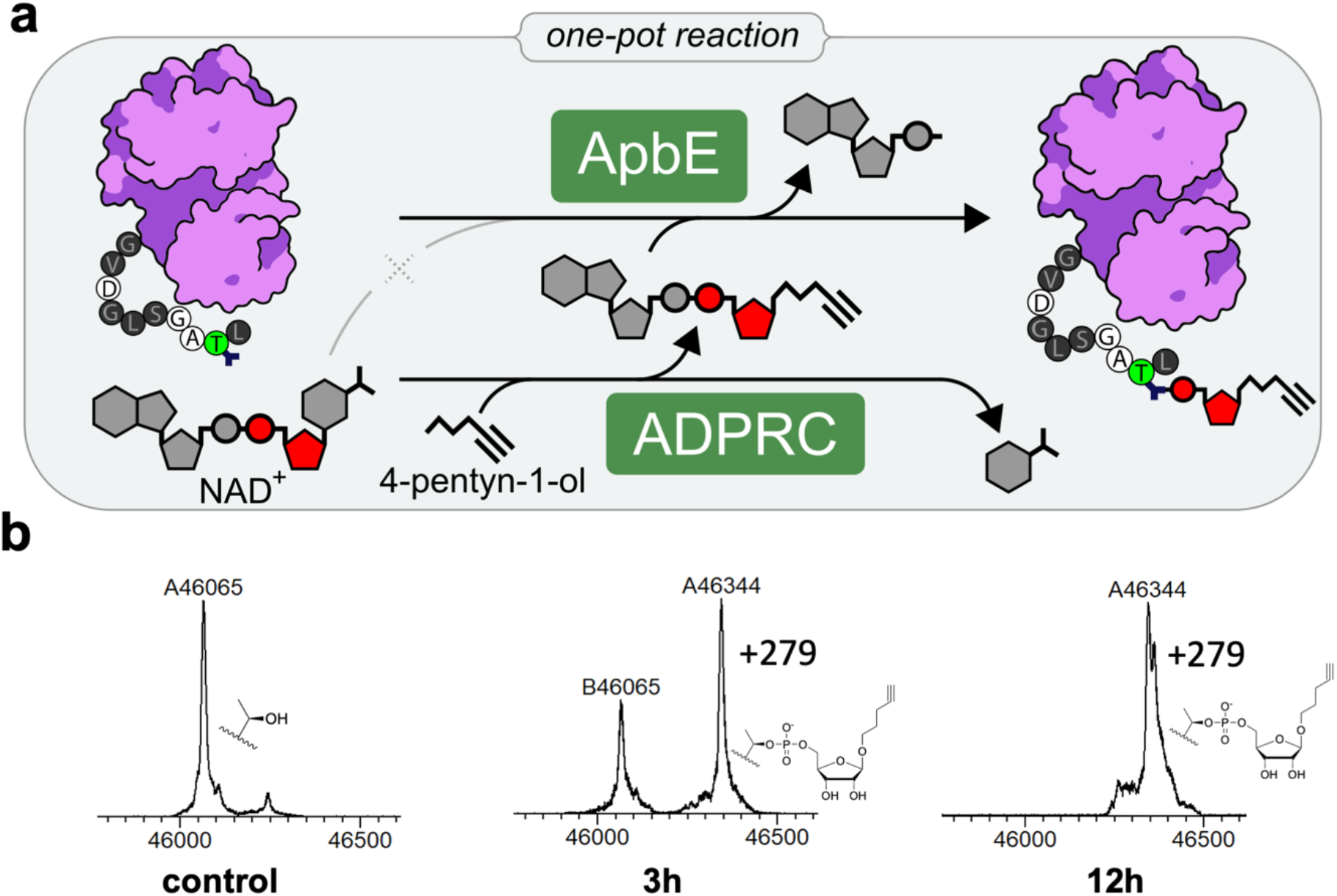
One-pot click chemistry handle attachment (alkynyl-phospho-ribose) to F1-MBP using ADPRC and ApbE in Tris·HCl pH 8 starting from NAD^+^ and 4-pentyn-1-ol. a) Reaction scheme. b) Complete labelling was observed after 12 hours reaction by MS.

### Attaching small molecules to azido and alkynyl proteins

With the target proteins modified with the bioorthogonal click chemistry handles in hand, we next sought to demonstrate a reliable and stable reactivity of ADD-tagged proteins in a broad variety of typical bioconjugation applications. A frequently desired reactivity in chemical biology is the site-specific installment of small molecule chemical probes, ranging from post-translational modification (mimics), over immobilization handles, to fluorophores. We therefore initially attempted to conduct the classic Cu(I)-catalyzed azide-alkyne cycloaddition (CuAAC)^[48]^ to conjugate small molecules useful for various applications. As a proof of concept, we reacted commercial azide variants of biotin or the cyanine dye Cy3 with the alkynylated proteins NqrC and F1-MBP. Using polyacrylamide gel electrophoresis (PAGE) for analysis, we could conveniently confirm the successful Cy3 conjugation manifesting as a fluorescent protein band (**Figures 4a, 4b and S7**), while biotin-PEG4 modification was visible as a band shift (**Figure 4c**). In-gel fluorescence also demonstrated that the introduced alkyne handle remains active in CuAAC conjugations of NqrC with an internal tag or MBP with an N-terminal tag, both converted in apparently quantitative yields within 20 mins and 10 mins, respectively.

**Figure 4.**
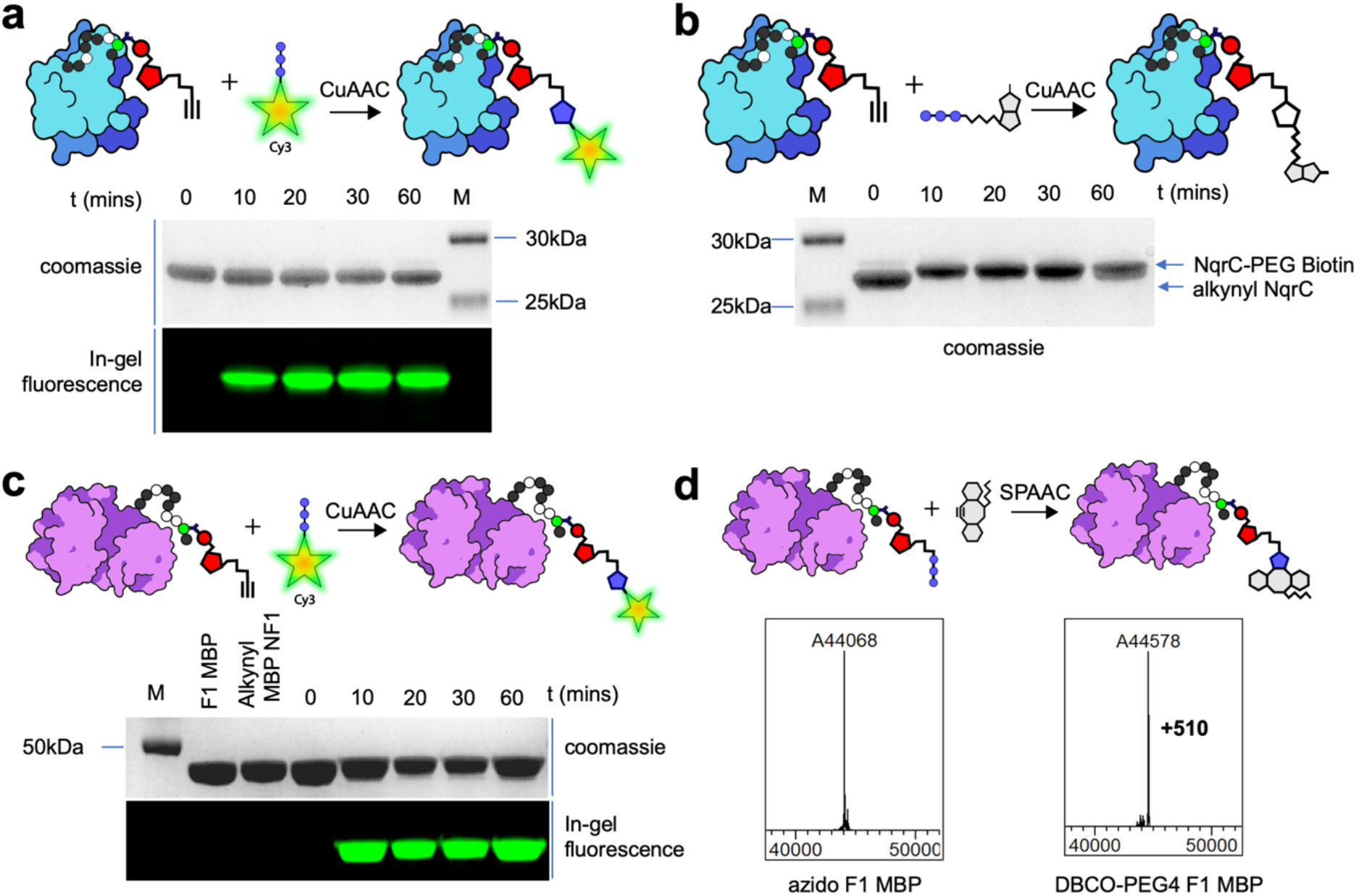
Small molecule decoration of modified target proteins using click chemistry. a) In-gel fluorescence was used to follow the reaction of alkynyl NqrC with Cy3 azide and showed saturation within 20 min. b) Reaction of alkynyl NqrC with azido-biotin-PEG4, demonstrating quantitative incorporation of biotin-PEG4 as indicated by a complete band shift after 10 minutes. c) Reactivity of N-terminally alkyne-modified MBP with Cy3 azide. d) SPAAC reaction of F1-MBP with DBCO PEG4 alcohol. MS analysis showed full conjugation as indicated by mass addition of 510 Da.

While CuAAC reactions are still widely employed, various adaptations have been developed over the years in an effort to increase the biocompatibility of click reactions. The strain-promoted azide-alkyne cycloaddition (SPAAC) is a popular alternative, prompting us to investigate its compatibility with our reactive protein handles.^[49,50]^ As a commercially available example molecule, we attempted to conjugate dibenzylcyclooctyne (DBCO)-PEG4 to alkynyl F1-MBP. Using MS, we confirmed the reactivity of the protein’s ADD-tagged azido handle, showing that a one-hour reaction resulted in full labelling (**Figure 4d**). In summary, these findings confirm that ADD-tags can be used to functionalize internal and terminally tagged proteins with useful small molecules in highly reactive CuAAC and SPAAC reactions in excellent yields.

### Biomolecular conjugations of proteins and DNA to ADD-tagged proteins

The ease to obtain clickable protein modifications and the excellent reactivity in the CuAAC reactions motivated us to next evaluate the possibility whether ADD-tagged proteins with complementary azido and alkynyl groups could be used in protein-protein coupling. Biorthogonal protein-protein conjugation is a powerful tool for late-stage biomolecule conjugation in therapeutics such as antibodies with fused domains,^[51]^ or protein constructs inaccessible by genetic fusions.^[28,30,52]^ Accordingly, we tested N-terminal-N-terminal (N-N), C-terminal-C-terminal (C-C), branched-branched (B-B), and N-terminal-branched (T-B) fusions. We used F1-MBP to assess N-N fusion, GBP-F2 and MBP-F1 to evaluate C-C fusion, NqrC to examine branched-branched fusion, and a combination of F1-MBP and NqrC to investigate N-terminal-branched fusions.

Following a procedure reported by Stengl et al., we used 10 µM equimolar concentrations of complementary functionalized proteins and 0.25 mM CuSO_4_ in aqueous buffer. Using SDS-PAGE analysis to monitor the formation of the protein coupling reaction over time, we observed successful formation of protein-protein conjugates with varying efficiencies (**Figure 5 and S8, Table S1**). The highest amount of homodimer formation was observed in MBP-MBP conjugates with N-N fusion, achieving approximately 25% conversion after 20 minutes of reaction time as estimated by densitometry (**Figure 5a**). Our C-C fusion model demonstrated a conjugation efficiency of approximately 10%, while both B-B and N-B fusions exhibited less than 5% conjugation (**Figure 5b, 5c, 5d**). To evaluate the functionality of the eGFP-protein conjugates, non-reducing SDS-PAGE gels were imaged at 473 nm excitation. As the fluorescence signals from both GFP-GFP and MBP-GFP conjugates were retained, we concluded that the conjugation process preserved the structural integrity and functionality of the fluorescent protein **(Figure S9)**. We hypothesize that the lower coupling efficiency is due to steric clashes between the proteins during conjugation. While several approaches could be employed to enhance efficiency, such as optimized reaction conditions,^[52,53]^ adjusted proteins concentrations,^[30]^ or linker engineering, we concluded that protein-protein coupling is in principle compatible with ADD-tagging. Recognizing that optimizations will likely be highly system-specific, we did not attempt to improve the reactions for our proof-of-concept setup further.

**Figure 5.**
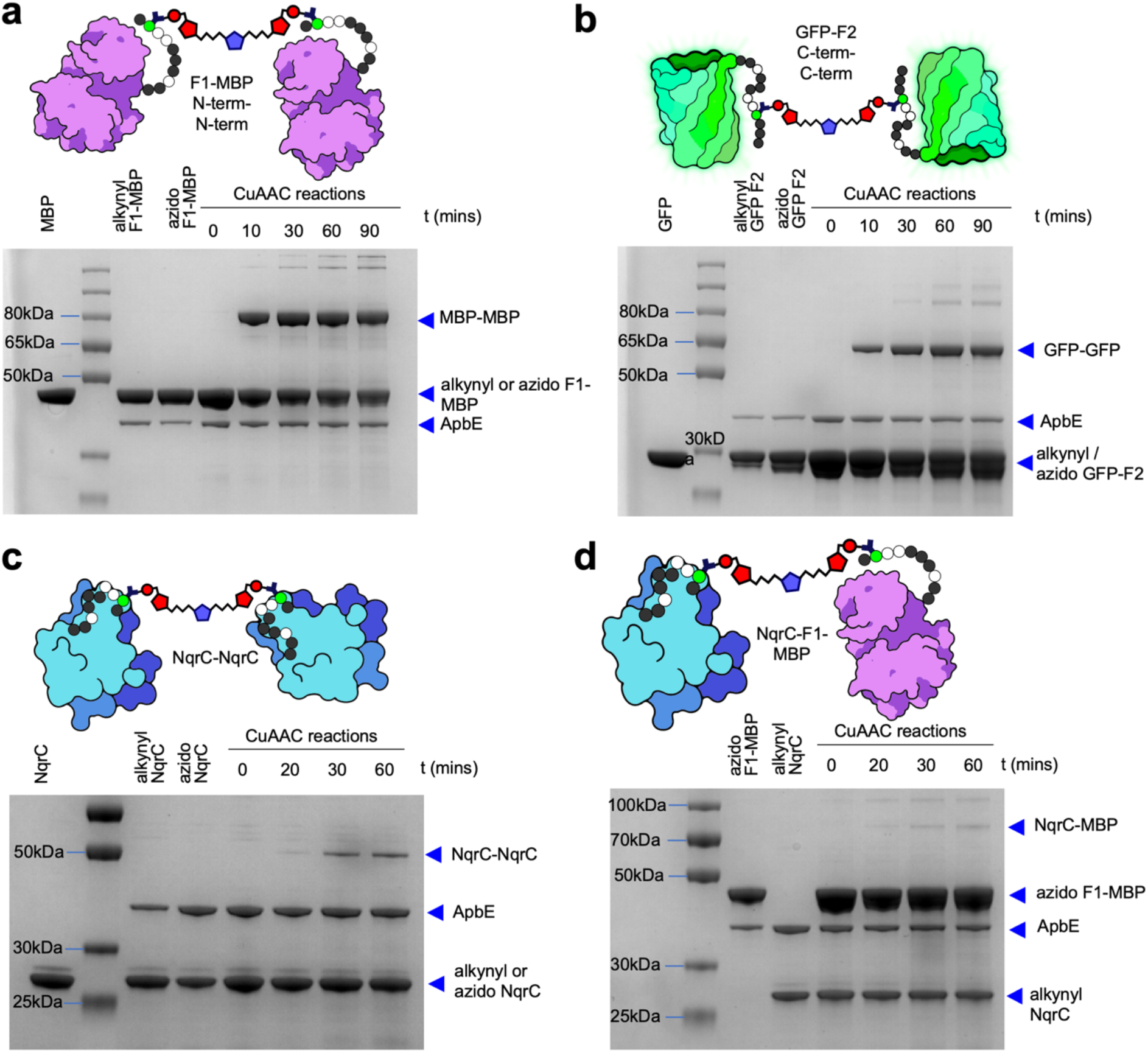
Protein-protein conjugation using CuAAC and ADD-tagged proteins. (a) Conjugation of terminal N-to-N fused proteins using complementary functionalized F1-MBP. (b) Conjugation of terminal C-to-C fused proteins involving alkynyl GFP-F2 and azido GFP-F2. (c) Conjugation of branched-branched proteins with complementary functionalized NqrC. (d) Conjugation of terminal-branched proteins involving NqrC and F1-MBP.

Instead, we were curious to investigate the capability of functionalized proteins to also form conjugates with nucleic acids. Access to such conjugates would broaden the potential applications of ADD-tag for applications such as DNA barcoding, optical tweezers, and material sciences.^[54,55]^ Using commercially synthesized azido DNA, we therefore conjugated alkynylated F1-MBP with a 45 bases long oligonucleotide. Using again SDS-PAGE to follow the reaction, we stained the gels with the highly sensitive SYBR Gold nucleic acid dye to detect the formation of the DNA-protein conjugate. Additionally, we hybridized the protein-ssDNA conjugate with Cy3-modified oligonucleotides, to visualize protein-dsDNA formation using in-gel fluorescence. The results indicated the successful, albeit low-yield formation of DNA-protein conjugates without any reaction optimization, as shown by SYBR Gold staining, in-gel fluorescence, and a band shift (**Figure 6**). We thus demonstrated that DNA-protein conjugates can be generated with alkynylated ADD-tags via CuAAC, with efficiency improvements likely possible via reaction engineering^[56]^ or using the SPAAC reaction^[57]^ to avoid Cu(II)-induced DNA degradation.^[58]^

**Figure 6.**
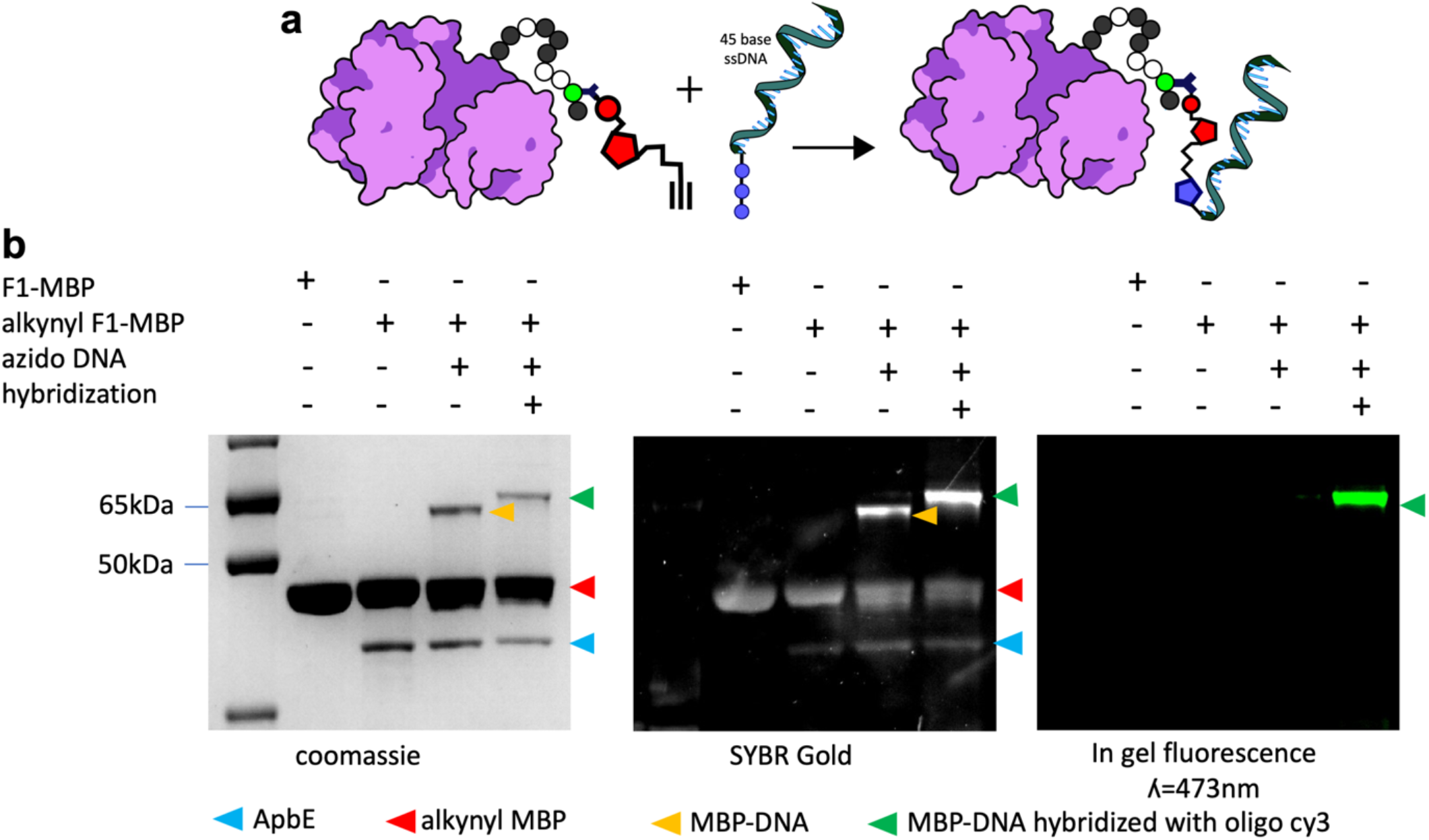
Demonstration of chemoenzymatic protein-DNA conjugation via ADD-tagged proteins. SDS-PAGE analysis reveals the creation of a protein-DNA conjugate, detected by Coomassie staining, SYBR Gold staining, and in-gel fluorescence analysis.

### Using ADD-tag for protein immobilization

As a final demonstration of the broad utility of ADDing and to expand the potential applications, we investigated the reactivity of modified proteins with azide agarose^[59]^ as a model for protein immobilization. For ease of detection, we selected alkynylated enhanced GFP (eGFP-F2). We conducted a 400 µl total reaction in a gravity flow column, using 200 µl of azide agarose and 50 µM alkynyl eGFP-F2, for 1 hour at room temperature, and found that GFP successfully reacted with the azide agarose, as evidenced by GFP immobilization on the column. This outcome was easily confirmed by both color and fluorescence detection, which remained even after extensive washing with an elution buffer containing 350 mM imidazole. In contrast, when we performed the same reaction using non-functionalized eGFP-F2 instead of the alkynylated protein as a control, GFP was nearly completely eluted (**Figure 7, S10**). We conclude that ADD-tagging can be used for protein immobilization and associated applications such as protein stabilization, biocatalyst recycling, advanced chromatography systems where proteins act as the stationary phase,^[59]^ or as a platform for ligand-antibody screening.^[60]^

**Figure 7.**
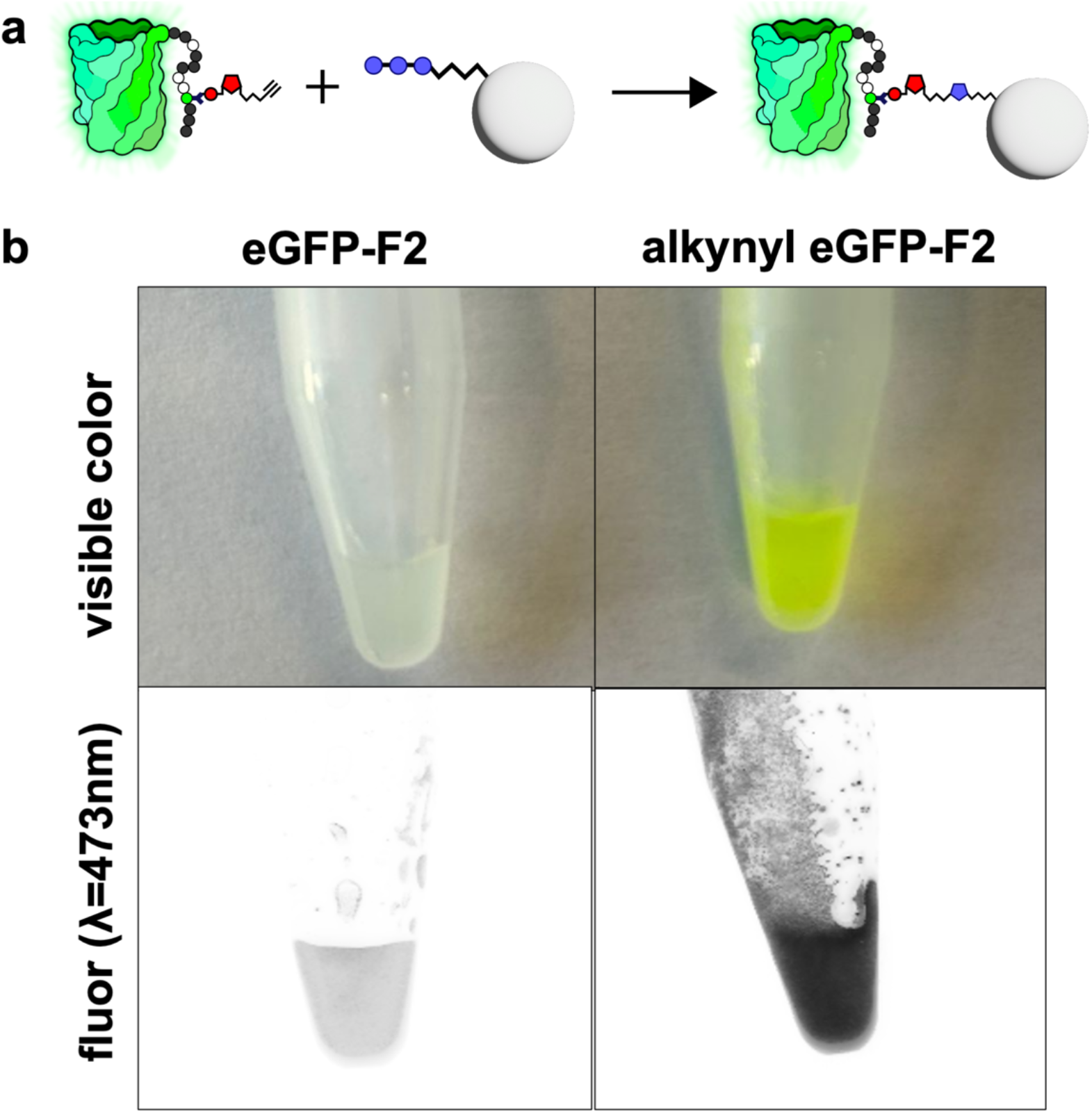
Immobilization of the EGFP model protein onto agarose azide. Covalent attachment is demonstrated by the retention of EGFP in the column after extensive washing with a buffer containing 350 mM imidazole.

## Conclusions

In summary, we present a streamlined enzymatic strategy for site-specific incorporation of bioorthogonal handles into proteins that we call “ADDing”. The small molecule donor substrates can be easily obtained via enzymatic ADPRC reactions, either separately, or together with bioconjugation in a one-pot, two-step method in combination with ApbE-mediated conjugation. Unlike other enzymatic labelling platforms, such as tubulin tyrosine ligase,^[23]^ lipoic acid ligase,^[61]^ or biotin ligase,^[26]^ which rely on synthetically modified substrates, our approach uses inexpensive small-molecule precursors and NAD^+^, offering advantages in cost, accessibility, and scalability.

The ADD tag’s minimal recognition motif (D-[GAIQ]-[IALVF]-[ST]-G-A-[ST]) consists of just seven semi-flexible amino acids, making it one of the shortest peptide tags for enzymatic labelling.^[47]^ This compact motif facilitates both internal and terminal labelling with minimal structural disruption, allowing for CuAAC- and SPAAC-based functionalization of small molecules, as well as the generation of protein-protein and protein-DNA assemblies. Additionally, compatibility with commercially available azide agarose enables efficient protein immobilization, supporting applications in affinity purification and enzyme display.

Although ADDing offers simple substrate synthesis and flexible recognition motifs, our method—like other tag-based systems—is not without limitations.^[17]^ It still requires relatively high substrate concentrations (1 mM in our case) and extended incubation times (3-12h) for efficient labelling.

However, we assume that (and currently explore if) the efficiency can be further improved through enzyme or reaction engineering. Additionally, the applicability of this method in vivo remains to be fully explored.

Overall, the minimal tag size and flexibility of the ADDing system make it well-suited for modular and orthogonal post-translational modifications. This platform not only expands the enzymatic conjugation toolkit but also offers significant promise for diverse applications in chemical biology and the development of biologics.

## References

[1] K. M. Hartung, E. M. Sletten, Chem 2023, 9, 2095–2109.

[2] M. S. Dillingham, M. I. Wallace, Org Biomol Chem 2008, 6, 3031.

[3] P. Giri, A. D. Pagar, M. D. Patil, H. Yun, Biotechnol Adv 2021, 53, 107868.

[4] Z. Fu, S. Li, S. Han, C. Shi, Y. Zhang, Signal Transduct Target Ther 2022, 7, 93.

[5] R. Liu, L. Liang, M. P. Lacerda, E. F. Freed, C. A. Eckert, New Frontiers and Applications of Synthetic Biology 2022, 147–158.

[6] K. Liu, M. Li, Y. Li, Y. Li, Z. Chen, Y. Tang, M. Yang, G. Deng, H. Liu, Mol Cancer 2024, 23, 62.

[7] A. Samantasinghar, N. P. Sunildutt, F. Ahmed, A. M. Soomro, A. R. C. Salih, P. Parihar, F. H. Memon, K. H. Kim, I. S. Kang, K. H. Choi, Biomedicine & Pharmacotherapy 2023, 161, 114408.

[8] M. J. Suskiewicz, BioEssays 2024, 46, e202300178.

[9] J. Hermann, L. Schurgers, V. Jankowski, Mol Aspects Med 2022, 86, 101066.

[10] A. K. Singh, S. Murmu, A. Krężel, ACS Omega 2022, 7, 46693–46701.

[11] E. A. Hoyt, P. M. S. D. Cal, B. L. Oliveira, G. J. L. Bernardes, Nat Rev Chem 2019, 3, 147–171.

[12] T. Pleiner, M. Bates, S. Trakhanov, C.-T. Lee, J. E. Schliep, H. Chug, M. Böhning, H. Stark, H. Urlaub, D. Görlich, Elife 2015, 4, e11349.

[13] R. V Chari, B. A. Martell, J. L. Gross, S. B. Cook, S. A. Shah, W. A. Blättler, S. J. McKenzie, V. S. Goldmacher, Cancer Res 1992, 52, 127–31.

[14] C. Uttamapinant, K. A. White, H. Baruah, S. Thompson, M. Fernández-Suárez, S. Puthenveetil, A. Y. Ting, PNAS. 2010, 107, 10914–10919.

[15] Y. Zhang, K.-Y. Park, K. F. Suazo, M. D. Distefano, Chem Soc Rev 2018, 47, 9106–9136.

[16] R. E. Bird, S. A. Lemmel, X. Yu, Q. A. Zhou, Bioconjug Chem 2021, 32, 2457–2479.

[17] J. Lotze, U. Reinhardt, O. Seitz, A. G. Beck-Sickinger, Mol Biosyst 2016, 12, 1731–1745.

[18] A. Amiri, S. Abedanzadeh, B. Davaeil, A. Shaabani, A. A. Moosavi-Movahedi, Q Rev Biophys 2024, 57, e6.

[19] K. Lang, J. W. Chin, ACS Chem Biol 2014, 9, 16–20.

[20] Y.-C. Wang, J. K. Dozier, L. S. Beese, M. D. Distefano, ACS Chem Biol 2014, 9, 1726–1735.

[21] S. A. Auger, S. Venkatachalapathy, K. F. G. Suazo, Y. Wang, A. W. Sarkis, K. Bernhagen, K. Justyna, J. V. Schaefer, J. W. Wollack, A. Plückthun, L. Li, M. D. Distefano, Bioconjug Chem 2024, 35, 922–933.

[22] M. Gerlach, T. Stoschek, H. Leonhardt, C. P. R. Hackenberger, D. Schumacher, J. Helma, 2019, pp. 327–355.

[23] D. Schumacher, J. Helma, F. A. Mann, G. Pichler, F. Natale, E. Krause, M. C. Cardoso, C. P. R. Hackenberger, H. Leonhardt, Angew. Chem. Int. Ed. 2015, 54, 13787–13791.

[24] D. Schumacher, H. Leonhardt, C. P. R. Hackenberger, J. Helma, 2019, pp. 167–189.

[25] D. Schumacher, O. Lemke, J. Helma, L. Gerszonowicz, V. Waller, T. Stoschek, P. M. Durkin, N. Budisa, H. Leonhardt, B. G. Keller, C. P. R. Hackenberger, Chem Sci 2017, 8, 3471–3478.

[26] S. A. Slavoff, I. Chen, Y.-A. Choi, A. Y. Ting, J Am Chem Soc 2008, 130, 1160–1162.

[27] I. Chen, M. Howarth, W. Lin, A. Y. Ting, Nat Methods 2005, 2, 99–104.

[28] M. D. Witte, J. J. Cragnolini, S. K. Dougan, N. C. Yoder, M. W. Popp, H. L. Ploegh, Proceedings of the National Academy of Sciences 2012, 109, 11993–11998.

[29] S. Yamazaki, Y. Matsuda, ChemistrySelect 2023, 8, e202302947.

[30] M. Baalmann, L. Neises, S. Bitsch, H. Schneider, L. Deweid, P. Werther, N. Ilkenhans, M. Wolfring, M. J. Ziegler, J. Wilhelm, H. Kolmar, R. Wombacher, Angew, Chem. Int. Ed. 2020, 59, 12885–12893.

[31] M. Fottner, J. Heimgärtner, M. Gantz, R. Mühlhofer, T. Nast-Kolb, K. Lang, J Am Chem Soc 2022, 144, 13118–13126.

[32] J. Tang, M. Hao, J. Liu, Y. Chen, G. Wufuer, J. Zhu, X. Zhang, T. Zheng, M. Fang, S. Zhang, T. Li, S. Ge, J. Zhang, N. Xia, Commun Chem 2024, 7, 87.

[33] F. B. H. Rehm, T. J. Tyler, Y. Zhou, Y.-H. Huang, C. K. Wang, N. Lawrence, D. J. Craik, T. Durek, Nat Chem 2024, 16, 1481–1489.

[34] J. Chatterjee, A. Bandyopadhyay, M. Pattabiraman, R. Sarkar, Chem. Commun. 2024, 60, 8978–8996.

[35] Y. Tong, M. Lee, J. Drenth, M. W. Fraaije, Bioconjug Chem 2021, 32, 1559–1563.

[36] Y. Tong, M. R. Loonstra, M. W. Fraaije, ChemBioChem 2022, 23, DOI 10.1002/cbic.202200144.

[37] A. V. Bogachev, A. A. Baykov, Y. V. Bertsova, Biochem Soc Trans 2018, 46, 1161–1169.

[38] Y. V Bertsova, M. V Serebryakova, V. A. Anashkin, A. A. Baykov, A. V Bogachev, FEMS Microbiol Lett 2019, 366, DOI 10.1093/femsle/fnz252.

[39] Y. V. Bertsova, M. S. Fadeeva, V. A. Kostyrko, M. V. Serebryakova, A. A. Baykov, A. V. Bogachev, Journal of Biological Chemistry 2013, 288, 14276–14286.

[40] T. F. Walseth, H. C. Lee, Biochimica et Biophysica Acta (BBA) - Molecular Cell Research 1993, 1178, 235–242.

[41] A. Kanavarioti, J. Lu, M. T. Rosenbach, T. Brian Hurley, Tetrahedron Lett 1991, 32, 6065–6068.

[42] F. Preugschat, G. H. Tomberlin, D. J. T. Porter, Arch Biochem Biophys 2008, 479, 114–120.

[43] H. Cahová, M.-L. Winz, K. Höfer, G. Nübel, A. Jäschke, Nature 2015, 519, 374–377.

[44] H. Hu, N. Flynn, H. Zhang, C. You, R. Hang, X. Wang, H. Zhong, Z. Chan, Y. Xia, X. Chen, PNAS. 2021, 118, e2025595118.

[45] R. Hofmann, G. Akimoto, T. G. Wucherpfennig, C. Zeymer, J. W. Bode, Nat Chem 2020, 12, 1008–1015.

[46] M. Gerlach, S. Schmitt, P. Cyprys, M.-A. Kasper, I. Mai, M. Klanova, A. Maiser, H. Leonhardt, C. P. R. Hackenberger, G. R. Fingerle-Rowson, A. M. Vogl, D. Schumacher, J. Helma, 2025, Biorxiv DOI 10.1101/2025.01.15.633119.

[47] Y. Tong, S. G. Kaya, S. Russo, H. J. Rozeboom, H. J. Wijma, M. W. Fraaije, J Am Chem Soc 2023, 145, 27140–27148.

[48] V. V. Rostovtsev, L. G. Green, V. V. Fokin, K. B. Sharpless, Angew. Chem. Int. Ed. 2002, 41, 2596–2599.

[49] X. Ning, J. Guo, M. A. Wolfert, G. Boons, Angew. Chem. Int. Ed. 2008, 47, 2253–2255.

[50] N. J. Agard, J. A. Prescher, C. R. Bertozzi, J Am Chem Soc 2004, 126, 15046–15047.

[51] M.-E. Goebeler, G. Stuhler, R. Bargou, Nat Rev Clin Oncol 2024, 21, 539–560.

[52] C. Kofoed, S. Riesenberg, J. Šmolíková, M. Meldal, S. Schoffelen, Bioconjug Chem 2019, 30, 1169–1174.

[53] A. Stengl, M. Gerlach, M.-A. Kasper, C. P. R. Hackenberger, H. Leonhardt, D. Schumacher, J. Helma, Org Biomol Chem 2019, 17, 4964–4969.

[54] G. Liszczak, T. W. Muir, Angew. Chem. Int. Ed. 2019, 58, 4144–4162.

[55] I. S. Gokulu, S. Banta, Trends Biotechnol 2023, 41, 575–585.

[56] J. Schwach, K. Kolobynina, K. Brandstetter, M. Gerlach, P. Ochtrop, J. Helma, C. P. R. Hackenberger, H. Harz, M. C. Cardoso, H. Leonhardt, A. Stengl, ChemBioChem 2021, 22, 1205–1209.

[57] S. L. Khatwani, J. S. Kang, D. G. Mullen, M. A. Hast, L. S. Beese, M. D. Distefano, T. A. Taton, Bioorg Med Chem 2012, 20, 4532–4539.

[58] C. J. Burrows, J. G. Muller, Chem Rev 1998, 98, 1109–1152.

[59] S. Punna, E. Kaltgrad, M. G. Finn, Bioconjug Chem 2005, 16, 1536–1541.

[60] M. S. Robescu, T. Bavaro, Molecules 2025, 30, 939.

[61] M. Fernández-Suárez, H. Baruah, L. Martínez-Hernández, K. T. Xie, J. M. Baskin, C. R. Bertozzi, A. Y. Ting, Nat Biotechnol 2007, 25, 1483–1487.

