## Supplementary material for "One-Pot Enzymatic ADDing of Click Chemistry Handles for Protein Immobilization and Bioconjugation of Small and Biomolecules": Materials and Methods, Supplementary Figures, Supplementary Tables

1. **SUPPLEMENTARY RESULTS**

**1.1. ApbE expression in *E. coli* BL21 AI and *riboflavin deficient* BL21 AI (∆ribB) strain.**

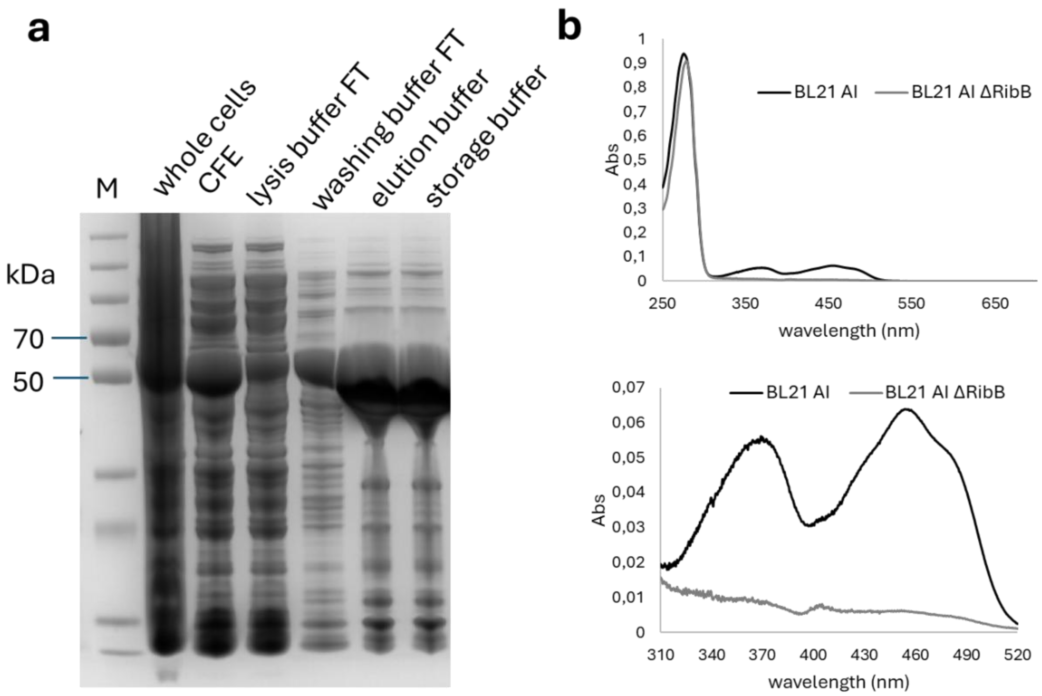

**Figure S1.** (a) SDS-PAGE analysis of ApbE expression in riboflavin deficient BL21 AI (∆*ribB*) strain and purification by Ni NTA gravity column. (b) Spectrum of ApbE purified from *E. coli* BL21 AI and *E. coli* BL21 AI (∆*ribB*) strains.

**1.2. ApbE conjugation reaction with ApbE produced in *E. coli* BL21 AI and *E. coli* BL21 AI (Δ*ribB*) strains.**

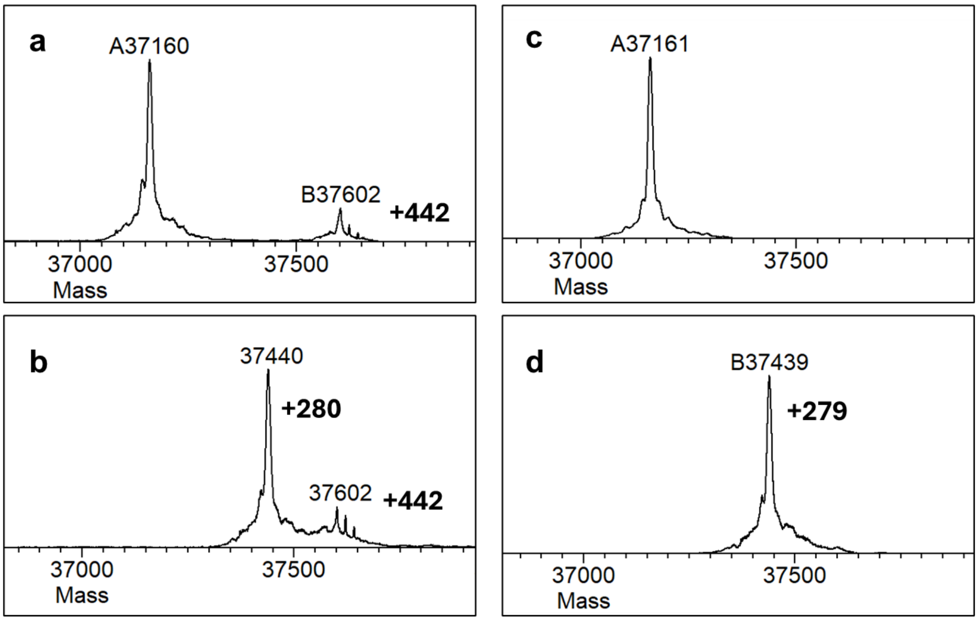

**Figure S2.** ApbE reactions with enzyme produced in *E. coli* BL21 AI (a,b) and *E. coli* BL21 AI (Δ*ribB*) (c,d). control reactions are shown in a and b, and reactions with alkynyl ADPR are shown in c and d. The detected mass of 37.602 Da indicates FMNylation caused by the presence of FAD in the purified ApbE with standard expression strains. The signal was not detected in reactions where the ApbE was purified from *E. coli* BL21 AI (Δ*ribB*).

#### 1.3. Optimization ADPRC reactions

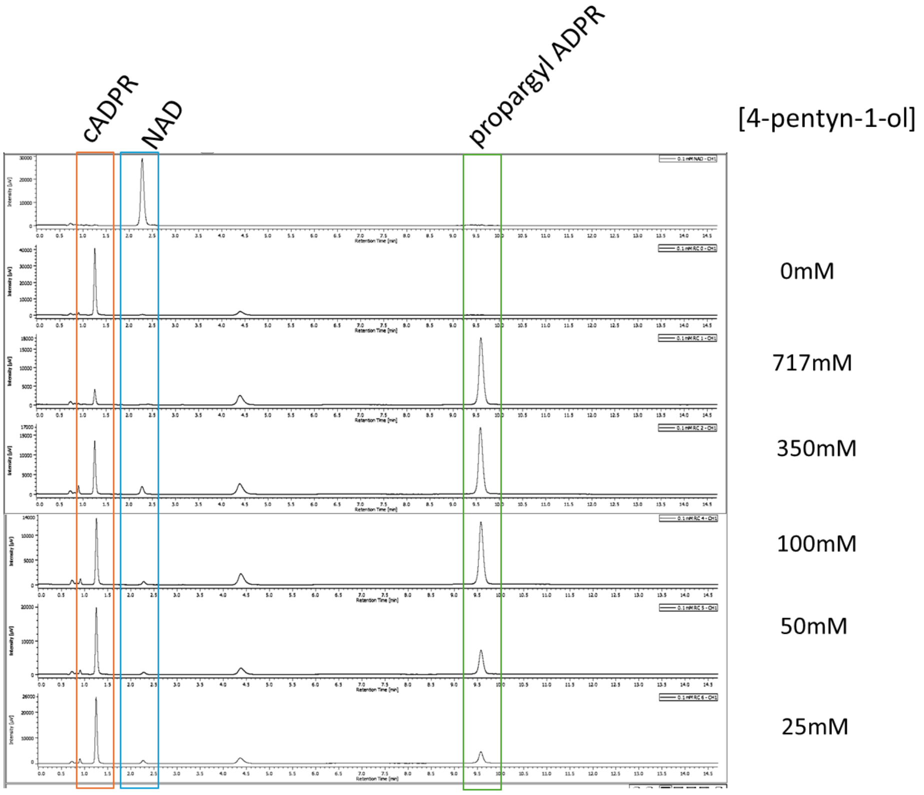

**Figure S3.** Optimization ADPRC reactions with various concentrations of 4-pentyn-1-ol.

#### 1.4. ApbE reaction with cADPR and NAD

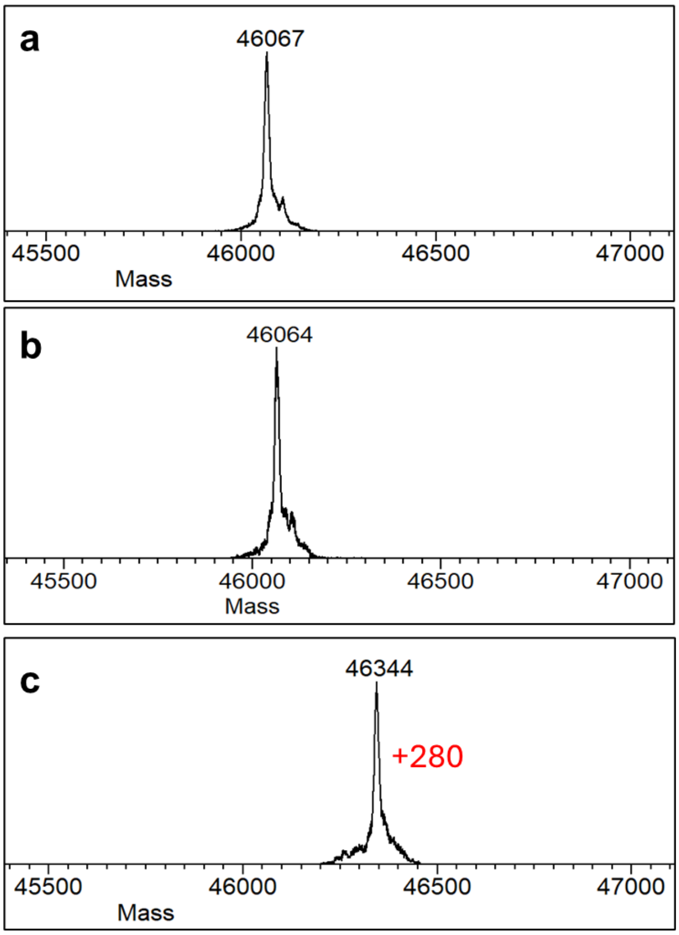

**Figure S4.** Intact MS analysis of ApbE reactions of MBP-NF1 with 1mM cADPR (a) 30mM NAD (b) and 1mM alkynyl ADPR (c). No increased mass was observed in the reactions with NAD and cADPR, indicating that neither of these compounds is a substrate for ApbE.

##### 1.5. ApbE reaction with GFP-F2 and SUMO iF1

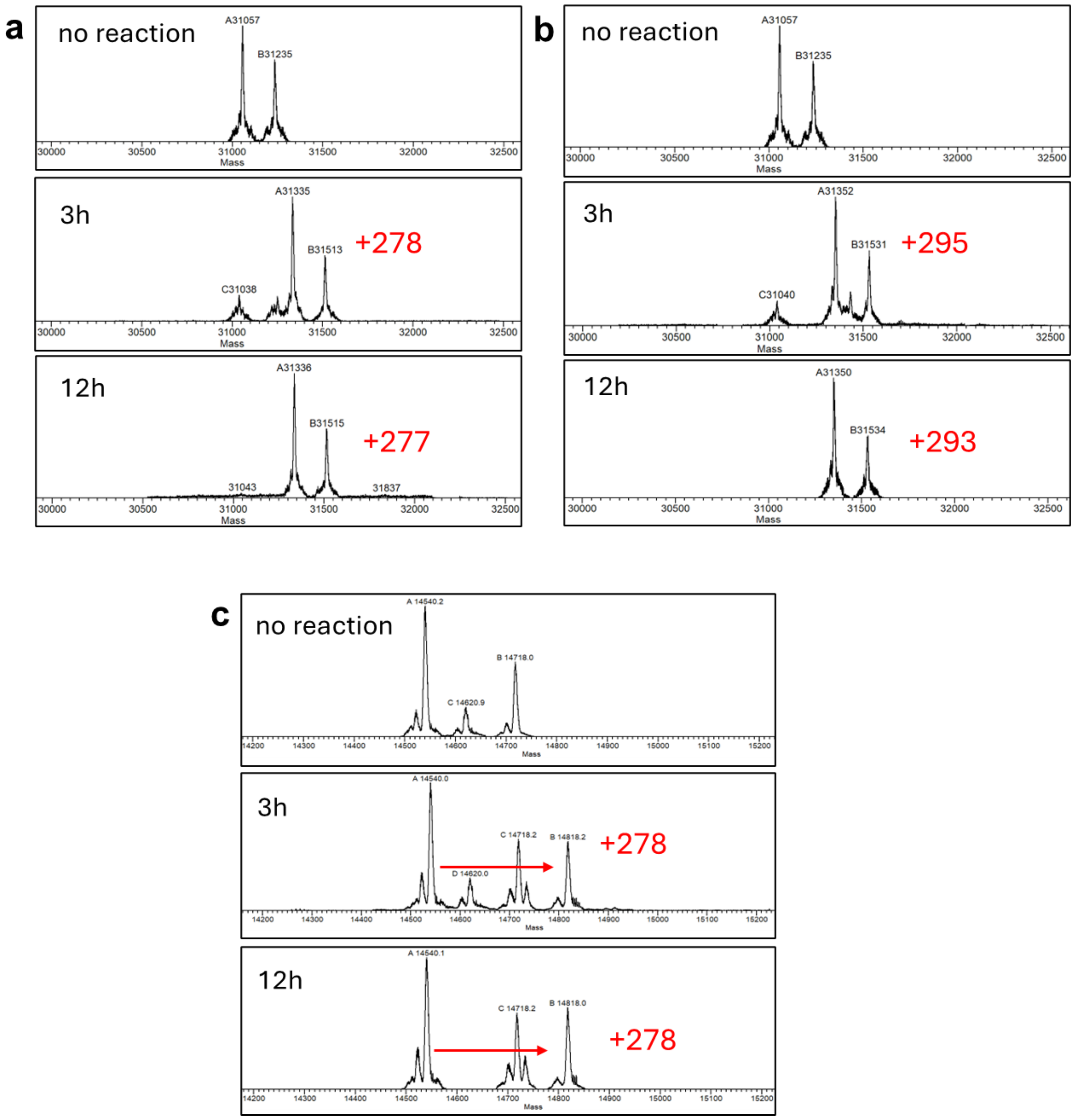

**Figure S5.** Intact MS analysis of EGFP-F2 (a = alkyne, b = azide) and SUMO iF1 (c) after reaction with ApbE for 3h and 12 h.

#### 1.6. Stability of threonyl-FMN linkage against various phosphatases and nucleases

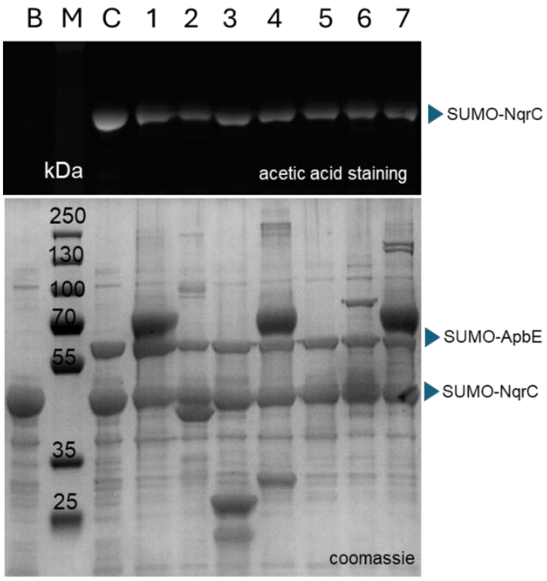

- B is blank, NqrC without ApbE;
- M is the protein ladder;
- 1 is treated with rSAP;
- 2 is treated with Antarctic phosphatase;
- 3 is treated with Lamda protein phosphatase;
- 4 is treated with NudC pyrophosphatase;
- 5 is treated with Nuclease P1;
- 6 is treated with DNAse 1;
- 7 is treated with RecBCD;

**Figure S6.** Stability of phosphodiester linkage between threonine and FMN against various phosphatases and nucleases.

**1.7. Full images of SDS-PAGE gel data presented in the Figure 4.**

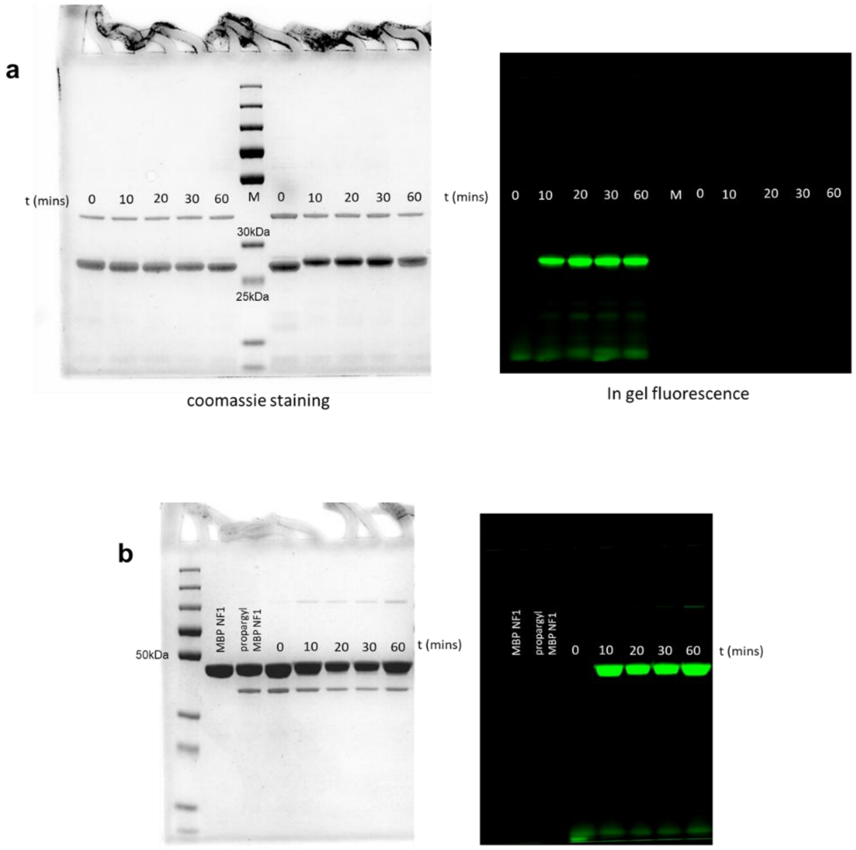

**Figure S7.** Full images of SDS-PAGE gel data presented in the Figure 4. (a) = figure 4a and 4c. (b) = figure 4b.

#### 1.8. C-C Protein-protein conjugation

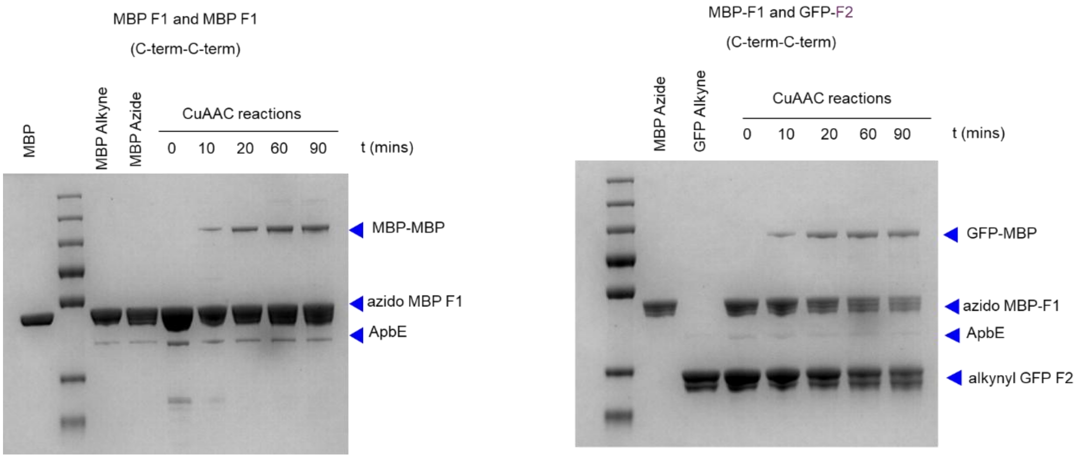

**Figure S8.** SDS-PAGE analysis of C-to-C conjugation of MBP CF1 and GFP CF2 and homodimer MBP CF1.

#### 1.9. GFP fluorescence detection of GFP-Protein conjugates.

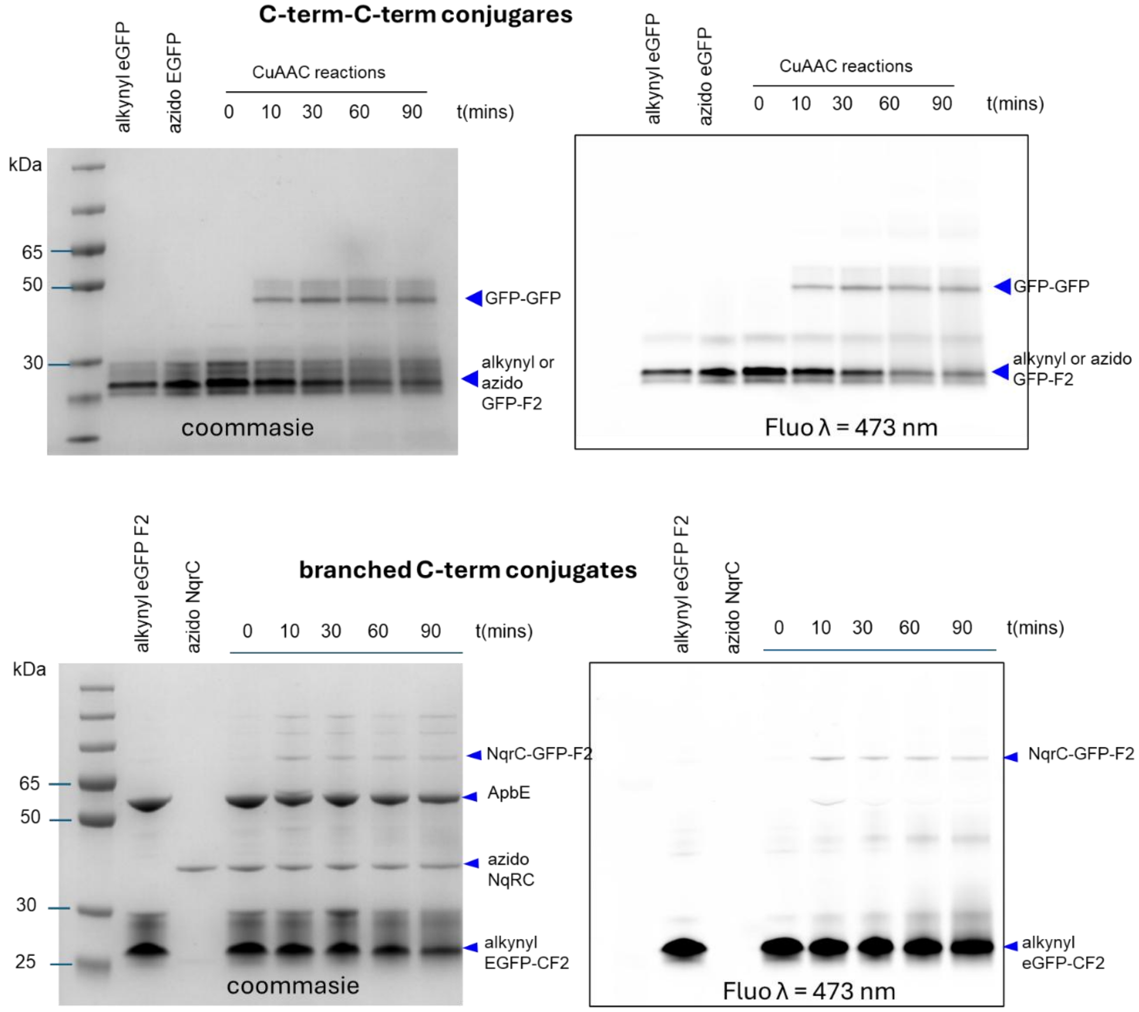

**Figure S9.** Fluorescence imaging of eGFP-protein conjugates on non-reducing SDS-PAGE gels.

##### 1.10. Efficiency of protein-protein conjugation

Table S1. Efficiency of protein-protein conjugation

| Protein 1 | Protein 2 | Efficiency |
|---|---|---|
| | **N-terminus – N-terminus** | |
| azido F1-MBP | alkynyl F1-MBP | ~25% |
| | **C-terminus – C-terminus** | |
| azido GFP-F2 | alkynyl GFP-F2 | ~10% |
| azido MBP-F1 | alkynyl MBP-F1 | ~5% |
| azido MBP-F1 | azido GFP-F2 | ~5% |
| | **Branched-Branched** | |
| azido NqrC | alkynyl NqrC | <5% |
| | **Branched-N-terminus** | |
| azido NqrC | alkynyl F1-MBP | <5% |
| | **Branched-C-terminus** | |
| azido NqrC | alkynyl F1-GFP | <5% |

#### 1.11. GFP CF2 immobilization in azido agarose

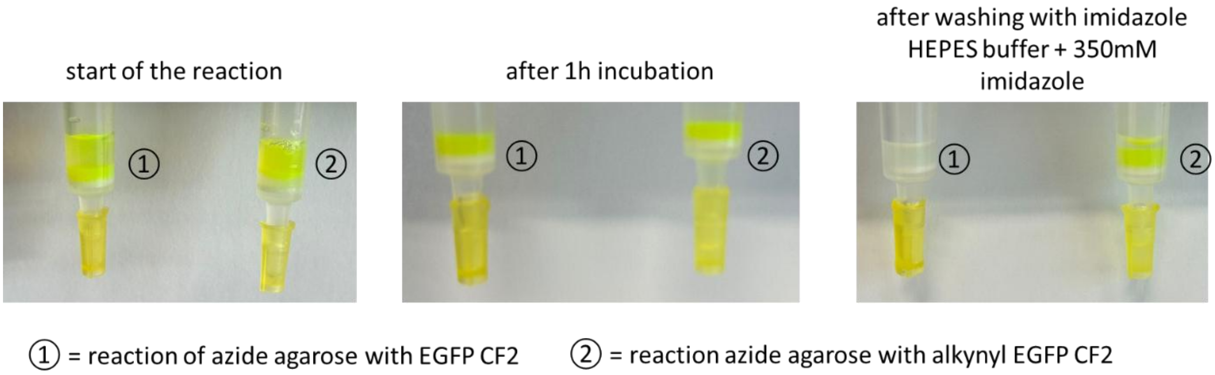

**Figure S10.** Step by step pictures of alkynylated model protein GFP CF2 immobilization in azide agarose.

##### 1.12. Amino acid sequence of ApbE and the target proteins.

**Table S2.** Amino acid sequence of the target proteins.

| Proteins | Amino acid sequence |
|---|---|
| ApbE | MGSSHHHHHHGSGLVPRGSASMSDSEVNQEAKPEVKPEVKPETHINLKVSDGS<br>SEIFFKIKKTTPLRRLMEAFAKRQGKEMDSLRFLYDGIRIQADQTPEDLDMEDNDI<br>IEAHREQIGGMEKPAEQVHLSGPTMGTTYNIKYIQQPGIADSKILQTEIDRLLEEV<br>NDQMSTYRKDSELSRFNQHTSSEPFAVSTQTLTVVKEAIRLNGLTEGALDVTVG<br>PLVNLWGFGPEARPDVVPTDEELNARRAITGIEHLTIEGNTLSKDIPELYVDLSTIA<br>KGWGVDVVADYLQSQGIENYMVEIGGEIRLKGLNRDGVPWRIAIEKPSVDQRSV<br>QEIIEPGDYAIATSGDYRNYFEQDGVRYSHIIDPTTGRPINNRVVSVTVLDKSCMT<br>ADGLATGLMVMGEERGMAVAEANQIPVLMIVKTDDGFKEYASSSFKPFLSK |
| NqrC | MGSSHHHHHHGSGLVPRGSASHRDLMSDSEVNQEAKPEVKPEVKPETHINLKV<br>SDGSSEIFFKIKKTTPLRRLMEAFAKRQGKEMDSLRFLYDGIRIQADQTPEDLDM<br>EDNDIIEAHREQIGGMKENAALDKQSKILQVAGIEAKGSKQIVELFNKSIEPRLVDF<br>NTGDFVEGDAANYDQRKAAKEASESIKLTAEQDKAKIQRRANVGVVYLVKDGDK<br>TSKVILPVHGNGLWSMMYAFVAVETDGNTVSGLTYYEQGETPGLGGEVENPAW<br>RAQWVGKKLFDENHKPAIKIVKGGAPQGSEHGVDGLSGATLTSNGVQNTFDFW<br>LGDMGFGPFLTKVRDGGLN |
| MBP-NF1 | MGSSHHHHHHGSGLVPRGSASGMGVDGLSGATLTSGGMKIEEGKLVIWINGDK<br>GYNGLAEVGKKFEKDTGIKVTVEHPDKLEEKFPQVAATGDGPDIIFWAHDRFGG<br>YAQSGLLAEITPDKAFQDKLYPFTWDAVRYNGKLIAYPIAVEALSLIYNKDLLPNP<br>PKTWEEIPALDKELKAKGKSALMFNLQEPYFTWPLIAADGGYAFKYENGKYDIKD<br>VGVDNAGAKAGLTFLVDLIKNKHMNADTDYSIAEAAFNKGETAMTINGPWAWSN<br>IDTSKVNYGVTVLPTFKGQPSKPFVGVLSAGINAASPNKELAKEFLENYLLTDEGL<br>EAVNKDKPLGAVALKSYEEELAKDPRIAATMENAQKGEIMPNIPQMSAFWYAVR<br>TAVINAASGRQTVDEALKDAQTNSSSNNNNNNNNNNLGIEGRI |
| MBP-CF1 | MGSSHHHHHHGSGLVPRGSASGMKIEEGKLVIWINGDKGYNGLAEVGKKFEKD<br>TGIKVTVEHPDKLEEKFPQVAATGDGPDIIFWAHDRFGGYAQSGLLAEITPDKAF<br>QDKLYPFTWDAVRYNGKLIAYPIAVEALSLIYNKDLLPNPPKTWEEIPALDKELKA<br>KGKSALMFNLQEPYFTWPLIAADGGYAFKYENGKYDIKDVGVDNAGAKAGLTFL<br>VDLIKNKHMNADTDYSIAEAAFNKGETAMTINGPWAWSNIDTSKVNYGVTVLPTF<br>KGQPSKPFVGVLSAGINAASPNKELAKEFLENYLLTDEGLEAVNKDKPLGAVALK<br>SYEEELAKDPRIAATMENAQKGEIMPNIPQMSAFWYAVRTAVINAASGRQTVDE<br>ALKDAQTNSSSNNNNNNNNNNLGIEGRIGGGVDGLSGATLTS |
| GFP-CF2 | MGSSHHHHHHGSGLVPRGSASGMVSKGEELFTGVVPILVELDGDVNGHKFSVS<br>GEGEGDATYGKLTLKFICTTGKLPVPWPTLVTTLTYGVQCFSRYPDHMKQHDFF<br>KSAMPEGYVQERTIFFKDDGNYKTRAEVKFEGDTLVNRIELKGIDFKEDGNILGH<br>KLEYNYNSHNVYIMADKQKNGIKVNFKIRHNIEDGSVQLADHYQQNTPIGDGPVL<br>LPDNHYLSTQSALSKDPNEKRDHMVLLEFVTAAGITLGMDELYKGIEENLYFQSG<br>VDGLSGATL |
| SUMO-iF1 | MGSSHHHHHHGSGLVPRGSASHMSDSEVNQEAKPEVKPEVKPETHINLKVSDG<br>SSEIFFKIKKTTPLRRLMEAFAKRQGKEMGGGVDGLSGATLTSGGDSLRFLYDGI<br>RIQADQTPEDLDMEDNDIIEAHREQI |

**Table S3.** Oligonucleotides used for DNA-protein conjugation.

| Oligos | Sequence (5'-3') |
|---|---|
| ssDNA 5'-Azide | /5AzideN/ATCACCGACTGCCCATAGAGAGGAAGGGAAGAGGAAGAAAGAG<br>GG |
| Comp_ssDNA_5'<br>-Cy3 | /5Cy3/CCCTCTTTCTTCCTCTTCCC |

#### 2. MATERIALS AND METHODS

##### 2.1. Plasmid Construction

The vectors pBAD SUMO ApbE, pBAD SUMO NqrC, and pET F1-MBP, pET MBP-F1, pET SUMO iF1 were obtained from previous studies all with 6x histidine tag in the N-terminus [1,2]. The construction of pBAD eGFP-F2 with 6x histidine tag in the N-terminus was created using Gibson assembly methods. The eGFP gene was amplified by PCR from an existing eGFP construct in our lab (addgene #185485), incorporating primer extensions for the F2 tag and the TEV cleavage site. All target protein sequences, and their corresponding tags are detailed in the Supplementary Information.

##### 2.2. Protein Expression and Purification

###### *apo* ApbE Expression and Purification

Purified ApbE, expressed in common strains like NEB10β and BL21 (DE3), binds tightly to FAD. To obtain apo ApbE, we expressed ApbE fused with a SUMO protein using a pBAD SUMO ApbE vector in a riboflavin-deficient *E. coli* strain, BL21 AI Δ*RibB*. An overnight preculture was inoculated into LB media containing 50 µg/ml ampicillin and 20 µg/ml riboflavin and cultured until the optical density (OD) reached 0.8. The cells were then washed twice to remove excess riboflavin from the medium. The riboflavin-free cells were resuspended in LB and induced with 0.02% arabinose. They were then incubated for 72 hours at 200 rpm and 17°C. For harvesting, the cells were centrifuged at 6000 rpm for 30 minutes.

ApbE was purified following previously established procedures. The cells were dissolved in 50 mM Tris-HCl (pH 7.5) with 0.5 mM PMSF and sonicated for 10 minutes (5 seconds on, 10 seconds off) to lyse the cells. Afterward, the mixture was centrifuged at 11,000 rpm for 60 minutes to obtain the supernatant. The supernatant was filtered through a 20 µm filter and then applied to a pre-incubated gravity Ni-NTA column. The column was rotated at 4°C for 30 minutes. Subsequently, the column was washed with 50 mM Tris-HCl (pH 7.5) containing 20 mM imidazole. ApbE was eluted in 50 mM Tris-HCl (pH 7.5) with 300 mM imidazole and desalted using a PD-10 desalting column by exchanging the buffer to 50 mM Tris-HCl (pH 7.5). The protein aliquots were then flash-frozen and stored at -80°C.

###### NqrC Expression and Purification

NqrC was expressed as a SUMO fusion protein from a pBAD SUMO NqrC construct in NEB10β cells. When the culture reached an optical density (OD) of 0.8, it was induced with 1 mM IPTG and incubated at 24°C for 24 hours. Bacterial cells were harvested by centrifugation at 6,000 rpm for 20 minutes, and protein purification was carried out using a protocol similar to that used for ApbE.

###### Expression and Purification of F1-MBP and MBP-F1

F1-MBP and MBP-F1 are maltose-binding protein (MBP) fusions that contain a C-terminal and N-terminal histidine tag, along with a 10-amino acid F1 tag (GVDGLSGATLTS). These proteins were expressed from a pET vector in E. coli BL21 DE3 cells. The cells were induced at an optical density (OD) of 0.8 with 1 mM IPTG and incubated at 24°C for 24 hours. After incubation, the bacterial cells were harvested by centrifugation at 6000 rpm for 20 minutes. Protein purification was then performed using a protocol similar to that of ApbE.

###### Expression and Purification eGFP F2

eGFP-F2 was expressed from the pET vector in the *E. coli* BL21 DE3 strain. The expression was carried out in TB media supplemented with 50 µg/ml ampicillin at 24°C and 200 rpm for 24 hours. Following this, the cells were harvested by centrifugation for 30 mins at 4500 rpm. To purify eGFP-F2, the cells were resuspended in 20 mM Tris-HCl (pH 8) with the addition of 0.5mM PMSF and 500mM NaCl. The cells were sonicated for 10 minutes (5 seconds on, 10 seconds off) to lyse the cell. Additional heat purification was done by incubating the sonicated cells in 60ºC for 30 mins and then centrifuged at 11,000 rpm for 60 minutes to obtain the supernatant. The supernatant was filtered using a 20 µm filter and applied to a pre-calibrated Ni-NTA gravity column, which was rotated at 4°C for 30 minutes. The column was subsequently washed with 20 mM Tris-HCl (pH 8), 500 mM NaCl, and 20 mM imidazole. eGFP-F2 was then eluted from the column using 20 mM Tris-HCl (pH 8), 500 mM NaCl, and 500 mM imidazole. The eluted protein was desalted with a PD-10 desalting column by exchanging the buffer to 20 mM Tris-HCl (pH 8) and 150 mM NaCl. Finally, protein aliquots were flash-frozen and stored at -80°C.

##### 2.3. Enzymatic Synthesis of pent-4-yn-1-yl ADP-ribose and 3-azido-propyl ADP-ribose

Synthesis of pent-4-yn-1-yl- and 3-azido-propyl ADP-ribose was carried out using A. californica ADP-ribosyl cyclase (ADPRC; Sigma-Aldrich) with NAD as the precursor. The reactions were prepared in a 50 mM HEPES buffer (pH 7.0) that was supplemented with 5 mM $MgCl_2$. Each reaction mixture contained 10 mM NAD, 2 µM A. californica ADP-ribosyl cyclase (ADPRC; Sigma-Aldrich), and 2 µL of either 4-pentyn-1-ol or 4-azido propanol as substrates. The incubations were performed at 37 °C for 30 minutes. The reaction was terminated by denaturing the ADPRC at 95 °C for 10 minutes. The mixtures were then centrifuged at 13,300 rpm for 10 minutes to pellet the denatured ADPRC. Finally, the reaction products were analyzed using ultra-high-performance liquid chromatography (UHPLC).

##### 2.4. UHPLC detection of propargyl ADP-ribose and azido ADP-ribose

The detection of propargyl ADP-ribose and azido ADP-ribose was conducted using UHPLC as described in previous method [3].

##### 2.5. Attachment of click chemistry handles to target proteins

The attachment of propargyl ribose phosphate and azido ribose phosphate to target proteins was catalyzed by ApbE in reactions ranging from 30 to 500 µL. These reactions contained 3 mg/mL target proteins, 1 mg/mL ApbE, 1 mM ADP-ribosyl alkyne or ADP-ribosyl azide, 1 mM EDTA, and 10 mM MgCl2, all in 50 mM Tris-HCl at pH 8, incubated at 30°C for 3 hours. Depending on the intended purpose, further purification of the target protein from ApbE was carried out using a His-tag column. To effectively purify the target proteins using this His-tag column, we utilized an untagged version of ApbE. To achieve this, the SUMO portion of the SUMO-ApbE fusion was cleaved by SUMO protease at a ratio of 1:5 (ApbE SUMO to SUMO protease) at 4°C overnight. The His-tagged target protein bound to the column while non-tagged ApbE was washed away. The purified protein was then eluted and desalted as described in the target protein purification section.

Before utilizing the proteins for the CuAAC reaction, any excess propargyl ADP-ribose or azido ADP-ribose was achieved by buffer exchange to PBS or HEPES buffer 4x using a 10 or 30 kDa Amicon spin column.

##### 2.6. ESI-MS detection of modified target proteins

To confirm the incorporation of click chemistry handles into target proteins, we used Electrospray Ionization Mass Spectrometry (ESI-MS) with a Waters® Xevo® G2 Tof system, which includes a quadrupole/time-of-flight (QToF) mass spectrometer and a PDA detector. Protein separation was achieved using an Acquity BEH C4 column (150 × 2.1 mm, 1.7 µm) maintained at 40°C. The mobile phase consisted of 0.1% formic acid in water (solvent A) and acetonitrile (solvent B) at a flow rate of 0.3 mL/min. The gradient program started at 5% solvent B, increased to 70% over 10 minutes, then ramped up to 95% B from 10 to 13 minutes, and finally returned to 5% B by 20 minutes for re-equilibration. Protein samples were diluted to a concentration of 1–5 µM and analyzed in ESI-positive ion mode using a 5µL injection volume. Charge density spectrum was processed and deconvoluted with MagTran software, which also generated the data visualizations displayed in Figures 2, 3, 4, S2, S4, and S5.

##### 2.7. CuAAC reaction with small molecules

CuAAC reactions with Cy3 Azide CuAAC reactions were performed using Cy3 azide with 20 µM alkynyl-MBP and 1 mM Cy3 azide (Jena Biosciences) or biotin PEG4 azide (Sigma). The reaction mixture also included 0.25 mM CuSO4, 1.25 mM THPTA (Tris(benzyltriazolylmethyl)amine), 5 mM aminoguanidine, and 5 mM sodium ascorbate, all dissolved in 1x HEPES pH 7 at room temperature[4]. Reactions were quenched at various time points by adding excess EDTA. Afterward, proteins were separated using SDS-PAGE and visualized through in-gel fluorescence detection on a Typhoon FLA 9500 Imager (λ=473). The proteins were subsequently stained with Coomassie blue and further analyzed using a Gel Doc Imager (Bio-Rad).

##### 2.8. SPAAC reaction with DBCO PEG alcohol

SPAAC reaction of azido MBP-NF1 with DBCO PEG alcohol was performed in 50 mM HEPES buffer pH 7 for 1h at 25ºC. The mixture contains 20 µM azido MBP-NF1 and 1 mM DBCO PEG alcohol. The excess DBCO PEG alcohol was washed with PBS and purified by Amicon ultra centrifugal kit (30 kDa cutoff) 5 times. The final reaction was analyzed using UPLC-MS.

##### 2.9. CuAAC for protein-DNA conjugation

CuAAC reactions were conducted using 20 µM alkynyl-MBP NF1 in combination with a fourfold excess of azide-DNA (IDT DNA), totaling a volume of 20 µM. The reactions took place in a 50 mM HEPES buffer at pH 7, containing 0.25 mM $CuSO_4$, 1.25 mM THPTA, 5 mM aminoguanidine, and 5 mM sodium ascorbate, at a temperature of 25°C for 1.5 hours. To quench the reactions, 100 mM EDTA was added immediately. In one sample, we hybridized the reaction with 20 bp Cy3 DNA oligonucleotides that were complementary to the 20 bp 3'-end of the azide DNA. The samples were then analyzed using SDS-PAGE and initially visualized with a Typhoon FLA 9500 (λ=473). Following this, the gels were stained with SYBR Gold and visualized using a UV filter.

##### 2.10. CuAAC for protein-protein conjugation

CuAAC reactions for protein-protein conjugation were performed using 20 µM of complementary functionalized target proteins. The reaction mixture included 0.25 mM $CuSO_4$, 1.25 mM THPTA, 5 mM aminoguanidine, and 5 mM sodium ascorbate in a 50 mM HEPES buffer at pH 7[5]. The reactions took place at room temperature with gentle shaking and were quenched at various time points by adding 125 mM EDTA. Protein samples were analyzed using SDS-PAGE and stained with Coomassie blue to estimate conversion efficiency through densitometric analysis.

##### 2.11. CuAAC for Protein Immobilization

A reaction mixture (200 µL) was prepared containing 100 µM propargyl GFP CF2, 0.5 mM $CuSO_4$, 2.5 mM THTPA, 10 mM aminoguanidine, and 10 mM sodium ascorbate in a 50 mM HEPES buffer at pH 7. For comparison, a control reaction was also conducted with 100 µM GFP CF2. The mixtures were then loaded into 200 µL azide agarose columns that had been pre-equilibrated with 50 mM HEPES buffer at pH 7. The reactions were carried out for 1.5 hours at room temperature. After the reaction, the columns were washed with the 50 mM HEPES buffer at pH 7, which contained 350 mM imidazole. The immobilized GFP remained in the column after washing and was visualized using the Typhoon imaging system FLA 9500 at a wavelength of 473 nm.

### References

[1] Y. Tong, M. Lee, J. Drenth, M. W. Fraaije, *Bioconjug Chem* **2021**, *32*, 1559–1563.

[2] Y. Tong, M. R. Loonstra, M. W. Fraaije, *ChemBioChem* **2022**, *23*, DOI 10.1002/cbic.202200144.

[3] J. Yoshino, S. Imai, **2013**, pp. 203–215.

[4] S. I. Presolski, V. P. Hong, M. G. Finn, *Curr Protoc Chem Biol* **2011**, *3*, 153–162.

[5] A. Stengl, M. Gerlach, M.-A. Kasper, C. P. R. Hackenberger, H. Leonhardt, D. Schumacher, J. Helma, *Org Biomol Chem* **2019**, *17*, 4964–4969.
